# Nutrient-Sensing Nuclear Receptor *PPARα* Controls Liver Proteostasis

**DOI:** 10.1101/2025.09.01.673563

**Authors:** Jinse Kim, Noor Bains, Kaylin G. Maddox, Julia G. Hwang, Jackson Q. Margolis, Jinyu Jiang, Francesca Fantini, Sung Yun Jung, Jong Min Choi, Naveed Ziari, Hussein Mohammed, Marc Hellerstein, Ana P. Arruda, Sungwoo Choi, Kang Ho Kim, David D. Moore

## Abstract

Peroxisome proliferator-activated receptor alpha (*PPARα*) is a nuclear receptor that orchestrates metabolic adaptation to fasting by regulating hepatic lipid and glucose metabolism. While these functions are well established, its role in protein homeostasis, another energy-intensive process, has remained unclear. Here, we performed multi-omics analyses in wild-type and *PPARα* knockout mice treated with the *PPARα* agonist fenofibrate to define its broader role in proteostasis. Interestingly, chronic *PPARα* activation suppressed hepatic secretory pathways, reducing secretome gene expression and circulating serum proteins, and downregulated genes involved in endoplasmic reticulum (ER) translocation, glycosylation, folding, and trafficking. Concurrently, *PPARα* activation enhanced proteasome activity, which was associated with selective induction of proteasome 26S subunit non-ATPase (PSMD) family members. In addition, hepatic protein synthesis was strongly attenuated. This was associated with an increase in the inhibitory phosphorylation of the eukaryotic translation initiation factor 2α (eIF2α), that in turn was linked to disrupted iron homeostasis. Together, these findings identify *PPARα* as a regulator of proteostasis, suppressing protein synthesis and secretion while promoting protein degradation pathways. Beyond its canonical role in lipid and glucose metabolism, we conclude that *PPARα* exerts an additional energy-conserving function by coordinating proteostasis, expanding our understanding of its systemic metabolic role.

## Introduction

Peroxisome proliferator-activated receptor alpha (*PPARα*) is a ligand-activated nuclear receptor that plays a central role in regulating lipid and energy metabolism. It is highly expressed in the liver, where it promotes key metabolic processes—including fatty acid oxidation, ketogenesis, lipid transport, and gluconeogenesis—to maintain systemic energy balance^1,2^. Under nutrient-deprived conditions, free fatty acids released through adipose tissue lipolysis act as natural ligands that activate *PPARα*^3^. Once activated, *PPARα* enhances peroxisomal and mitochondrial fatty acid oxidation, stimulates gluconeogenesis and ketone body production, and modulates lipoprotein metabolism to maintain systemic energy homeostasis^4,5,6,7,8^. In clinical settings, fibrate drugs, synthetic *PPARα* agonists, are widely used to lower triglyceride levels and increase HDL cholesterol, particularly in patients with dyslipidemia^9,10^. These physiological functions and therapeutic applications highlight the essential role of *PPARα* in maintaining lipid and energy homeostasis. Our previous work showed that *PPARα* promotes energy conservation through direct activation of autophagy, but the broader role of *PPARα* in other aspects of hepatic energy balance remains unclear.

The proteostasis network is a finely tuned system that regulates proteins from their synthesis to degradation. Secreted and membrane proteins are folded in the endoplasmic reticulum (ER), while other proteins are translated and folded in the cytosol. Misfolded proteins are directed toward degradation pathways, primarily through autophagy or the ubiquitin–proteasome system. In contrast, properly folded secretory proteins are transported through the Golgi apparatus for further processing. This network contributes to energy balance in response to physiological conditions and prevents accumulation of toxic protein aggregates. Disruption of proteostasis by pathological conditions is associated with diseases including neurodegenerative disorders and type 2 diabetes^19,20^. Therefore, maintaining whole-body proteostasis is crucial, particularly in the liver, which is a central hub for both protein synthesis and degradation. Hepatocytes are the primary site for the production of circulating plasma proteins, including albumin, clotting factors, transport proteins, and components of the immune system^21^. In parallel, the liver actively participates in protein turnover, degrading intracellular and extracellular proteins to liberate amino acids that are used for the resynthesis of plasma proteins, channeled into gluconeogenesis to maintain blood glucose levels during nutrient-deprived conditions, or exported to support protein synthesis in peripheral tissues such as muscle, brain, and immune cells^22^. In light of emerging evidence suggesting a role for *PPARα* in protein metabolism and the maintenance of protein energy balance, we explored its function in hepatic proteostasis.

## Materials and Methods

### Animal Studies

All animal experiments were conducted in accordance with the ethical guidelines and regulations established by the University of California, Berkeley, Animal Care and Use Committee (Protocol ID: AUP-2021-12-14887). Animals were housed in an Association for Assessment and Accreditation of Laboratory Animal Care (AAALAC)-certified facility under the supervision of licensed technicians. Throughout the study, animal health and welfare were monitored daily by veterinary staff from the Office of Laboratory Animal Care (OLAC) to ensure compliance with established ethical and welfare standards. For all experiments, male mice were used. Wild-type C57BL/6J (JAX, stock # 000664) and *PPARα* knockout (KO) (JAX, stock # 008154) mice were obtained from the Jackson Laboratory. Animals were maintained on a standard chow diet (LabDiet 5053) under a 12-hour light/dark cycle with ad libitum access to food and water until they reached 8–12 weeks of age. Animals aged between 8-12 weeks old were fed either a chow diet (Envigo, TD.110180) or a 0.2% fenofibrate diet (Envigo, TD.160663) for two weeks, then sacrificed at 10:00 AM for analysis. To assess protein synthesis rates using the SUnSET assay, animals were intraperitoneally injected with puromycin at a dose of 40 nmol/g body weight. After 45 minutes, mice were sacrificed, and their organs were harvested, snap-frozen in liquid nitrogen, and stored at −80°C until further analysis.

### Bulk RNA-seq analysis

Hepatic gene expression in fenofibrate-fed WT and *PPARα* KO was profiled by bulk RNA sequencing conducted by Arraystar Inc. Briefly, total RNAs were extracted from snap-frozen liver tissue and purified using the RNeasy MinElute Cleanup Kit (Qiagen). Messenger RNA was enriched by oligo dT magnetic beads. Libraries were prepared with the KAPA Stranded RNA-seq Kits (Roche), and sequenced on the NovaSeq 6000 instrument (Illumina). Raw sequencing data were processed using the Galaxy platform (https://usegalaxy.org). Reads were aligned to the GRCm38/mm10 mouse genome using HISAT2, and gene counts were summarized with featureCounts. Differential gene expression analysis was performed using the Limma-Voom tool. Gene Ontology (GO) and Kyoto Encyclopedia of Genes and Genomes (KEGG) analyses were performed to identify *PPARα*-regulated targets. All raw sequencing data generated in this study have been deposited into the Sequence Read Archive (SRA) at the National Center for Biotechnology Information (NCBI) in the National Institutes of Health (NIH). The BioProject number is PRJNA1222824 and a temporary link during the review process is available at https://dataview.ncbi.nlm.nih.gov/object/PRJNA1222824?reviewer=8qn2lljqdsss6hrn3icqb40btc.

### Mass spectrometry-based proteomics

To profile the whole proteome, liver homogenates and precleared serum were used. Frozen liver tissue was homogenized by cryogenic grinding and lysed in ammonium bicarbonate lysis buffer (50 mM NH5CO3 and 1 mM CaCl2) with repeated freeze-thaw cycles. In serum samples, albumin and immunoglobulins were depleted using Proteome Purify 2 Mouse Immunodepletion resin (MIDR002, RnD Systems). Extracted proteins were trypsinized and peptides were recovered using an acetonitrile/formic acid solution. Lyophilized peptides were dissolved in loading solution (5% methanol and 0.1% formic acid) and analyzed using a nano-LC1000 system coupled to a Thermo Q-Exactive mass spectrometer (Thermo Fisher Scientific). Relative expression was quantified using the intensity-based absolute quantification (iBAQ) algorithm and normalized to the intensity-based fraction of the total (iFOT). Detailed procedures have been previously described (PMID:29718219 and 33168190).

### RNA extraction and quantitative RT-PCR analysis

Total RNA was extracted using TRIzol reagent (Thermo Fisher Scientific, Cat# 15596) according to the manufacturer’s protocol. Complementary DNA (cDNA) was synthesized from the isolated RNA using the qScript cDNA SuperMix (QuantaBio, Cat# 95048). Gene expression levels were assessed by quantitative real-time PCR (qRT-PCR) using the LightCycler 480 Real-Time PCR System (Roche) in combination with KAPA SYBR FAST qPCR Kits (KAPA Biosystems). Relative mRNA expression was quantified using the comparative cycle threshold (ΔΔCt) method, with Cyclophilin mRNA serving as the internal reference for normalization. Primer sequences are provided in material section.

### Immunoblot analysis

Mouse tissues were homogenized in RIPA lysis buffer (Thermo Fisher Scientific, Cat# 37576) supplemented with protease and phosphatase inhibitors (Thermo Scientific, Cat# PI78441). Protein concentrations were quantified using the Pierce BCA Assay Kit (Thermo Fisher Scientific, Cat# 23227). A total of 20–30 μg of protein was separated on either 8% or 4–12% NuPAGE Bis-Tris Protein Gels (Thermo Fisher Scientific, Cat# WG1002BOX or NP0322BOX) and transferred onto Immuno-Blot polyvinylidene difluoride (PVDF) membranes (Bio-Rad, Cat# 162-0177). Membranes were blocked with 2.5% milk in 1× TBST and incubated overnight at 4°C with primary antibodies, as detailed in 2.5.1 material section. The following day, membranes were washed with 1× TBST and incubated with appropriate secondary antibodies. Protein bands were detected using ECL reagents (Thermo Scientific, Cat# 34580 or 34095) and imaged using the iBright 1500 Imaging System (Thermo Fisher Scientific). Band intensity was normalized to a loading control and quantified using ImageJ software.

### Serum and Tissue Iron Quantification

Total iron levels in serum and liver tissue were quantified using the Iron Assay Kit (Sigma-Aldrich, Cat# MAK472) following the manufacturer’s instructions. Frozen serum samples were thawed prior to analysis, and 50 μL of serum was added to each well of a 96-well plate. The working reagent provided in the kit was subsequently added, mixed thoroughly, and incubated at RT for 40 minutes. OD was measured at 590 nm, and iron concentrations were calculated using the manufacturer’s equation. For tissue iron quantification, frozen liver samples (50 mg) were homogenized in 0.5 mL of RIPA lysis buffer, and the lysate was centrifuged to obtain the supernatant. Iron quantification in tissue samples was performed using the same assay and protocol as described for serum samples.

### Liver Heme Quantification

Liver heme levels were quantified using a Heme Assay Kit (Sigma-Aldrich, Cat# MAK316) according to the manufacturer’s instructions. Briefly, 50 mg of liver tissue was homogenized in RIPA buffer, followed by centrifugation to collect the supernatant. A 50 μL of the lysate was added to each well of a 96-well plate, and the heme reagent was subsequently added to each sample. Following a 5 mins incubation, absorbance was measured at 400 nm. Heme concentrations were calculated using the equation provided by the manufacturer.

### Proteasome assay

Liver tissue samples (50 mg) were homogenized in RIPA buffer and centrifuged, and the supernatant was used to assess proteasome activity. The proteasome activity assay was performed using the Proteasome Activity Assay Kit (Abcam, Cat# ab107921) according to the manufacturer’s instructions. Briefly, protein lysates were added to each well, followed by the addition of either a proteasome inhibitor or assay buffer. Proteasome substrate was then added to all samples, and the initial OD values were recorded. After incubation at 37°C for 30 minutes, the final OD values were measured to calculate proteasome activity.

### Heavy water labeling

Mice received an IP injection priming dose of 100% D₂O (Sigma-Aldrich 151882-250G) to rapidly attain an approximate 5% body water enrichment at 35 μl/g body weight. This enrichment level was subsequently maintained by providing the animals with 8% D₂O in their daily drinking water for 3 days. After 3days, mice were sacrificed, and the liver was harvested and homogenized in homogenization buffer (PBS containing 1 mM PMSF, 5 mM EDTA, and 1× Halt Protease Inhibitor Cocktail). The homogenized liver was then sonicated, followed by centrifugation at 10,000 × g for 10 minutes at 4°C. The resulting supernatant was collected and quantified using a BCA assay. 500 µg of protein samples were denatured and reduced by adding ammonium bicarbonate (ABC) solution, trifluoroethanol (TFE), and dithiothreitol (DTT) solution, followed by vortexing and incubation at 60°C for 1 hour. Samples were cooled to room temperature, alkylated with iodoacetamide (IAM) solution, and incubated for 1 hour in the dark. Excess IAM was quenched with DTT solution, incubated for 20 minutes in the dark. The samples were diluted with deionized water and ABC solution to achieve a final TFE concentration <5% and a pH of ∼8. Trypsin was added at a 1:50 (trypsin:protein) ratio, with trypsin prepared at 0.1 µg/µL in 25 mM ABC. Digestion was carried out at 37°C overnight. Reactions were terminated by adding neat formic acid. Samples were concentrated, centrifuged at 10,000 × g for 10 minutes, and the upper 90% of the supernatant was transferred to an LC-MS vial. The remaining sample was dried completely and reconstituted in LC-MS submission buffer. Peptides were analyzed via liquid chromatography-mass spectrometry (LC-MS).

### EM sample preparation and TEM image acquisition

Sample preparation for TEM and FIB-SEM was performed as previously described (A). In brief, mouse livers were perfused via the portal vein with saline, followed by fixation using a solution containing 2.5% glutaraldehyde and 2.5% paraformaldehyde in 0.1 M sodium cacodylate buffer (pH 7.4) (Electron Microscopy Sciences, catalog no. 15949). The liver tissues were then sectioned into 300-μm slices using a Compresstome and further incubated in a secondary fixative composed of 1.25% formaldehyde, 2.5% glutaraldehyde, 0.03% picric acid, and 0.05 M cacodylate buffer. For TEM imaging, samples were embedded in resin, sectioned into ultrathin slices using a Reichert Ultracut-S microtome, and examined using a JEOL 1200EX transmission electron microscope operated at 80 kV.

### Statistical Analysis

All animal data are presented as mean ± SEM. For comparisons involving two groups across two independent variables, two-way ANOVA followed by Tukey’s post hoc test was performed. Statistical significance was defined as p < 0.05.

### Antibodies

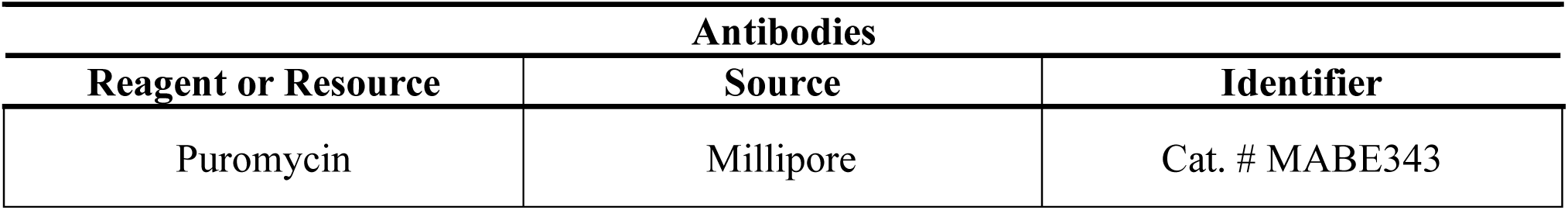
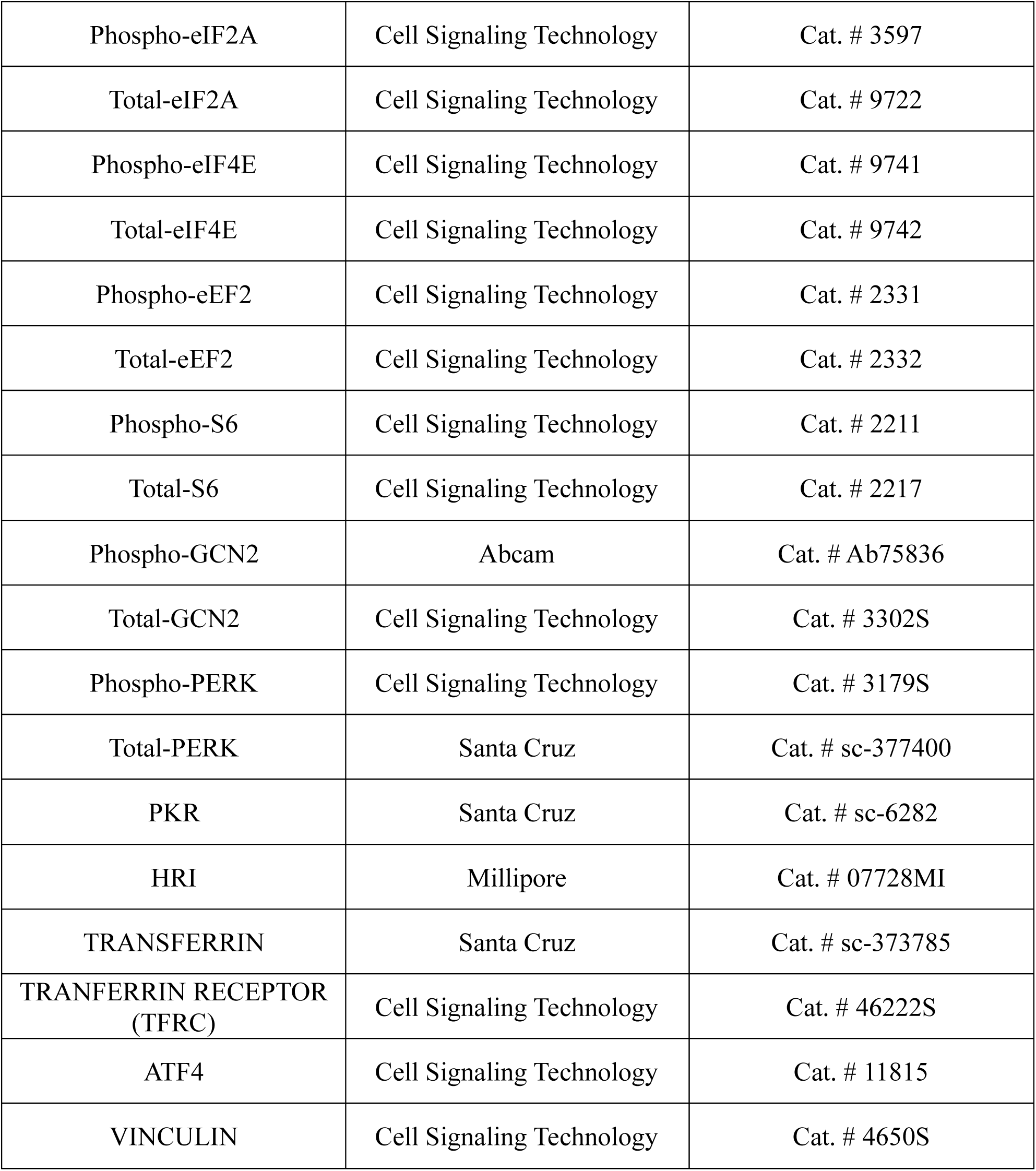

### qPCR Primers

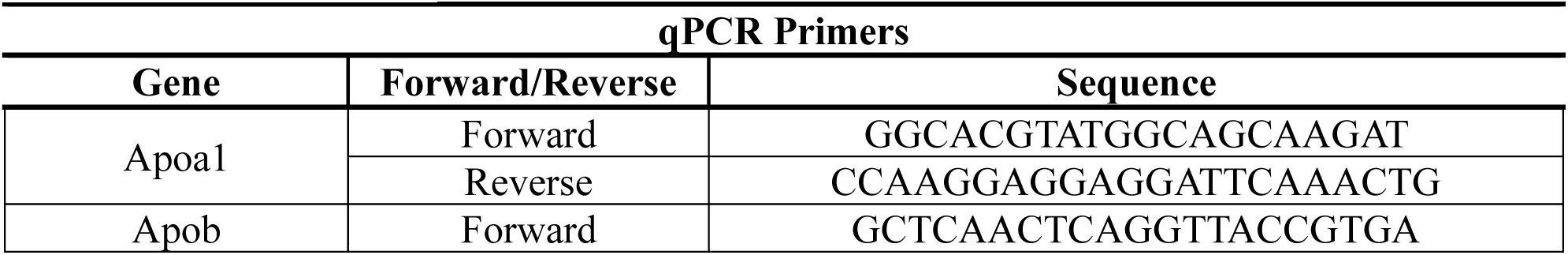
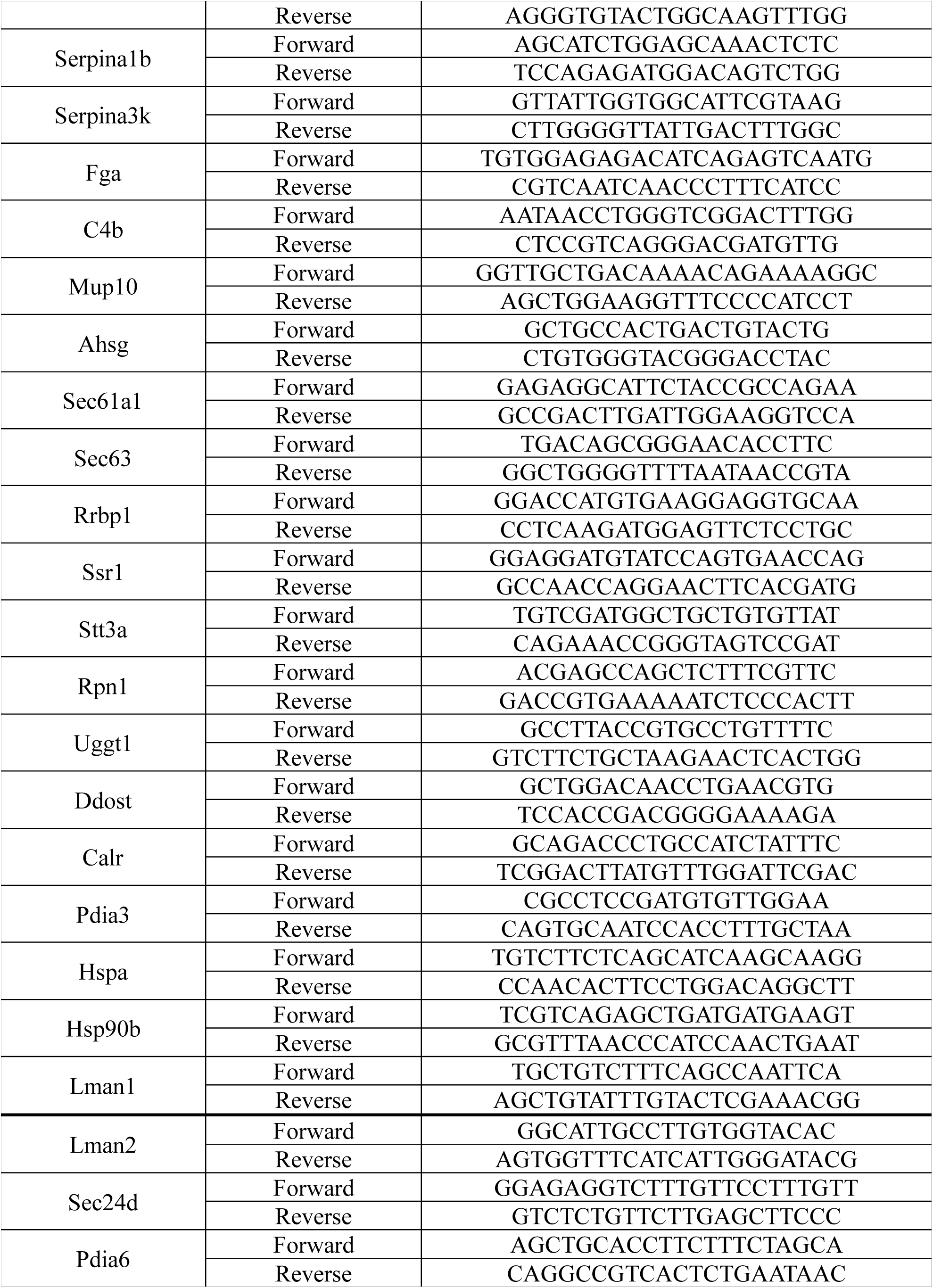
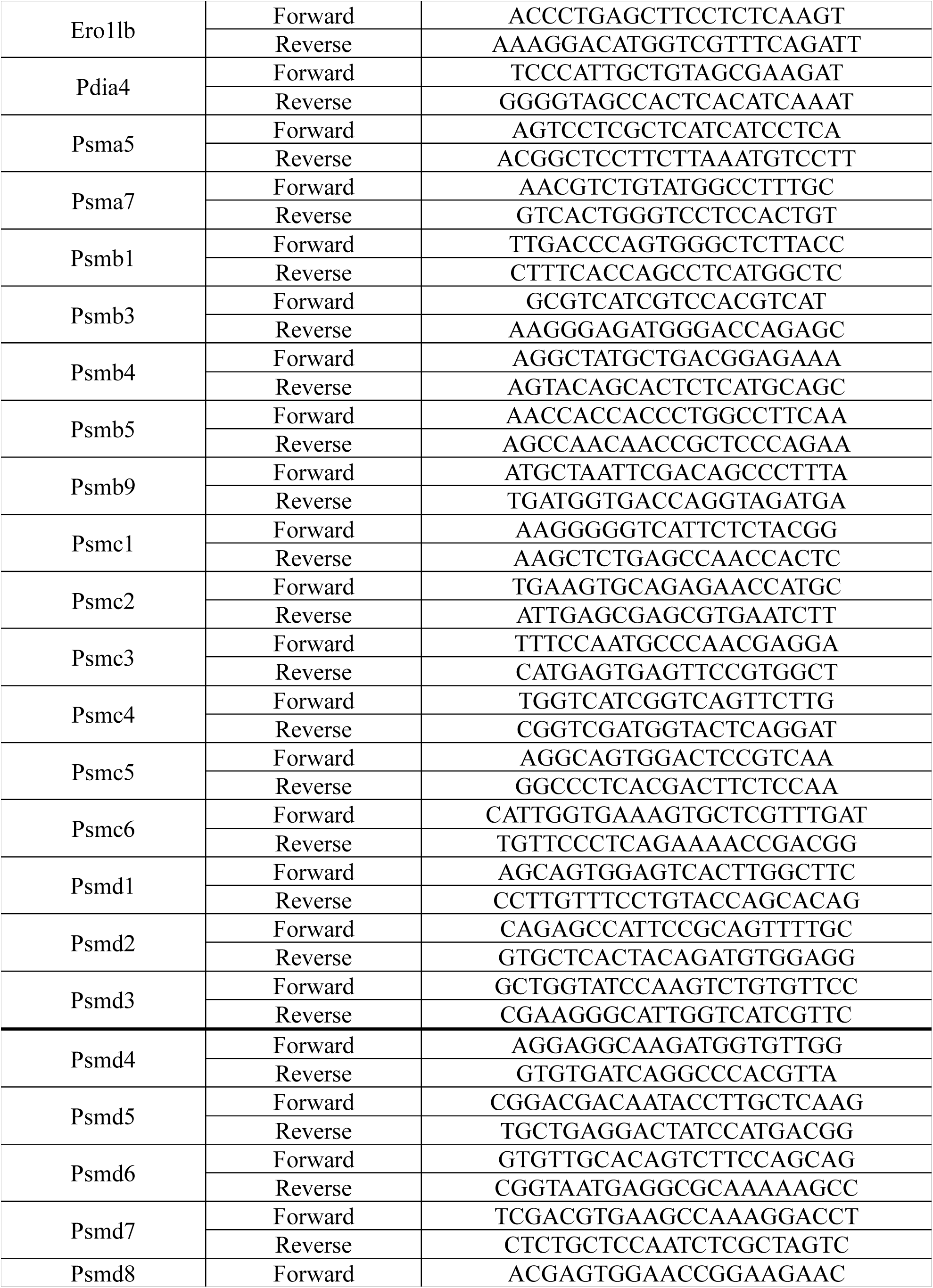
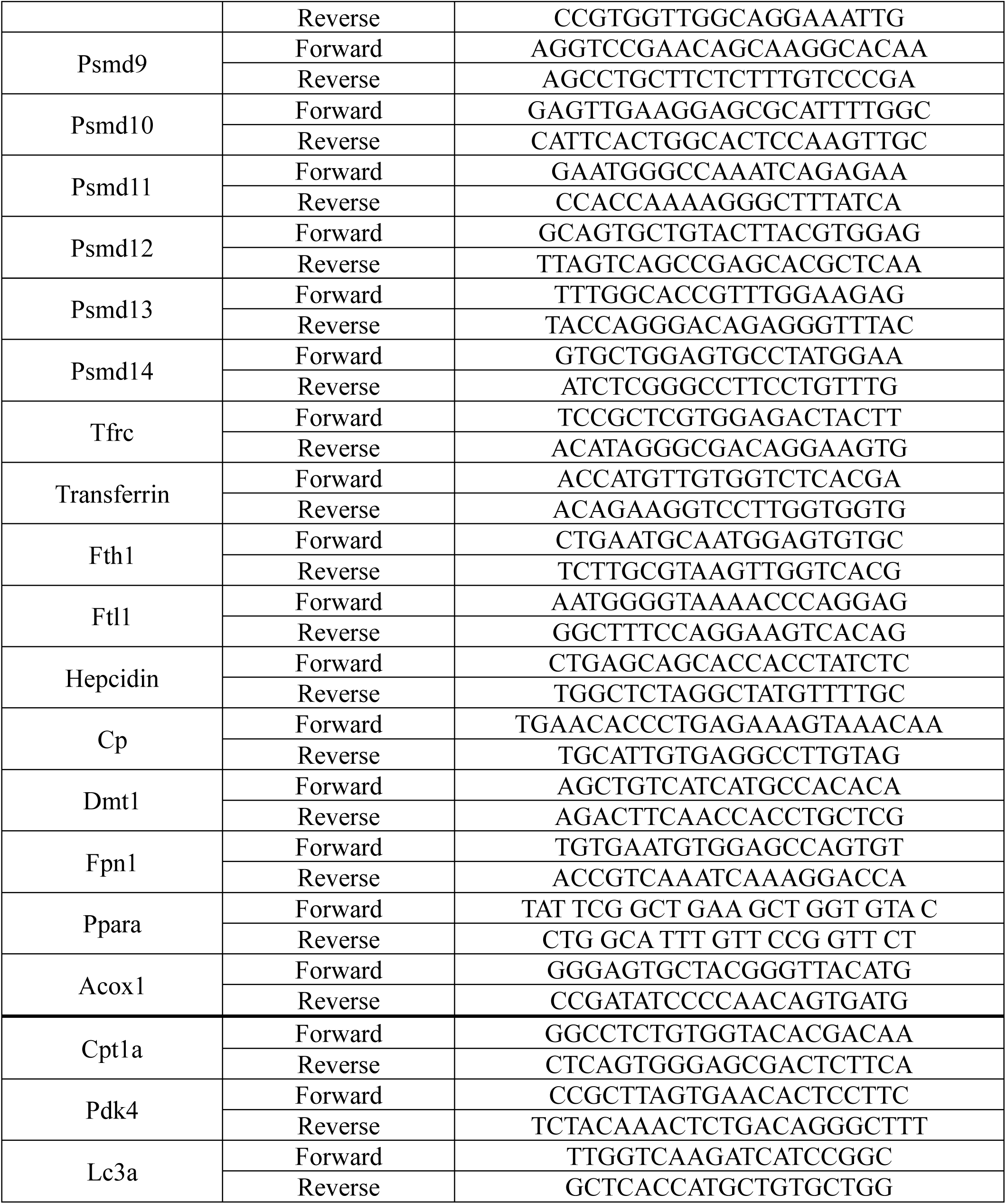

## Results

### Integrative Multi-Omics Analysis Reveals *PPARα*-Mediated Regulation of Hepatic Transcriptome, Proteome, and Post-Translational Modifications

To investigate the broader functions of peroxisome proliferator-activated receptor alpha (*PPARα*) beyond its known roles in coagulation and autophagy, wild-type (WT) and *PPARα* knockout (PKO) male mice were fed either a standard chow diet or a 0.2% fenofibrate-supplemented diet for two weeks. Liver samples were then collected for transcriptomic and proteomic analyses (Fig. 1). RNA-seq analysis identified 27,179 transcripts, of which 3,553 (13.1%) were significantly altered in fenofibrate-fed WT livers compared to untreated controls (P < 0.01). Among these, 1,999 transcripts (56.3%) were upregulated, while 1,554 (43.7%) were downregulated following fenofibrate treatment. Proteomic analysis detected 3,167 proteins, with 1,327 (41.9%) exhibiting significant changes in fenofibrate-fed WT livers (P < 0.01). Among these differentially expressed proteins, 375 (28.3%) were upregulated, whereas 952 (71.7%) were downregulated (Supplemental Fig. 1). Heatmap analyses of RNA-seq and proteomic data revealed distinct gene expression patterns in fenofibrate-fed WT livers compared to other groups (Fig. 1*A* and *B*). Additionally, principal component analysis (PCA) plots further confirmed that fenofibrate-fed WT liver formed a distinct cluster, highlighting the significant transcriptomic and proteomic alterations induced by *PPARα* activation (Fig. 1*C* and *D*). To gain a comprehensive understanding of the molecular mechanisms underlying *PPARα* activation, we conducted an integrative multi-omics analysis of RNA-seq and proteomic datasets from chow- and fenofibrate-fed WT mouse livers. This analysis identified a common set of 3,097 genes/proteins detected in both datasets. Among these, 1,211 transcripts (39.1%) and 1,327 proteins (42.8%) were significantly altered, with 658 genes exhibiting differential expression at both transcript and protein levels (Supplemental Fig. 1). The Pearson correlation coefficient for the 3,097 genes detected in both RNA-seq and proteomic datasets revealed a moderate positive correlation (*r* = 0.473598) between transcript and protein expression in chow-fed and fenofibrate-fed WT livers (Fig. 1*E*). In contrast, the Pearson correlation coefficient for the 658 differentially expressed genes at both transcript and protein levels indicated a strong positive correlation (*r* = 0.722504, Fig. 1*F*). Among the total 3,097 genes, 2,439 exhibited significant changes in expression at either the transcript or protein level. Notably, the majority of these genes were predominantly distributed along the y-axis, representing protein expression, rather than the x-axis, representing RNA expression. This pattern suggests the presence of post-translational regulation, indicating that protein abundance is influenced beyond transcriptional control. Next, we performed Gene Ontology (GO) enrichment analysis on the 658 differentially expressed genes at both transcript and protein levels to investigate the biological functions associated with *PPARα* activation (Supplemental Fig. 1*A* and *B*). As expected, the upregulated gene sets were significantly enriched in GO categories related to fatty acid metabolism, mitochondria-peroxisome function, and the PPAR signaling pathway, aligning with the well-established role of *PPARα* in lipid metabolism. In contrast, the downregulated gene sets were enriched in pathways associated with coagulation, amino acid biosynthesis, extracellular exosome, ER function, and iron binding. While the repression of specificcoagulation-related genes by *PPARα* was previously reported, the observed downregulation of pathways linked to amino acid metabolism, extracellular exosome, ER function, and iron homeostasis was unexpected, suggesting additional, previously unrecognized roles of *PPARα* activation in hepatic physiology (Supplemental Fig. 2*A* and *B*). Together, our integrative analysis of RNA-seq and proteomic data reveals the distinct molecular characteristics of *PPARα*-activated livers, uncovering unexpected targets of *PPARα*.

**Figure 1.**
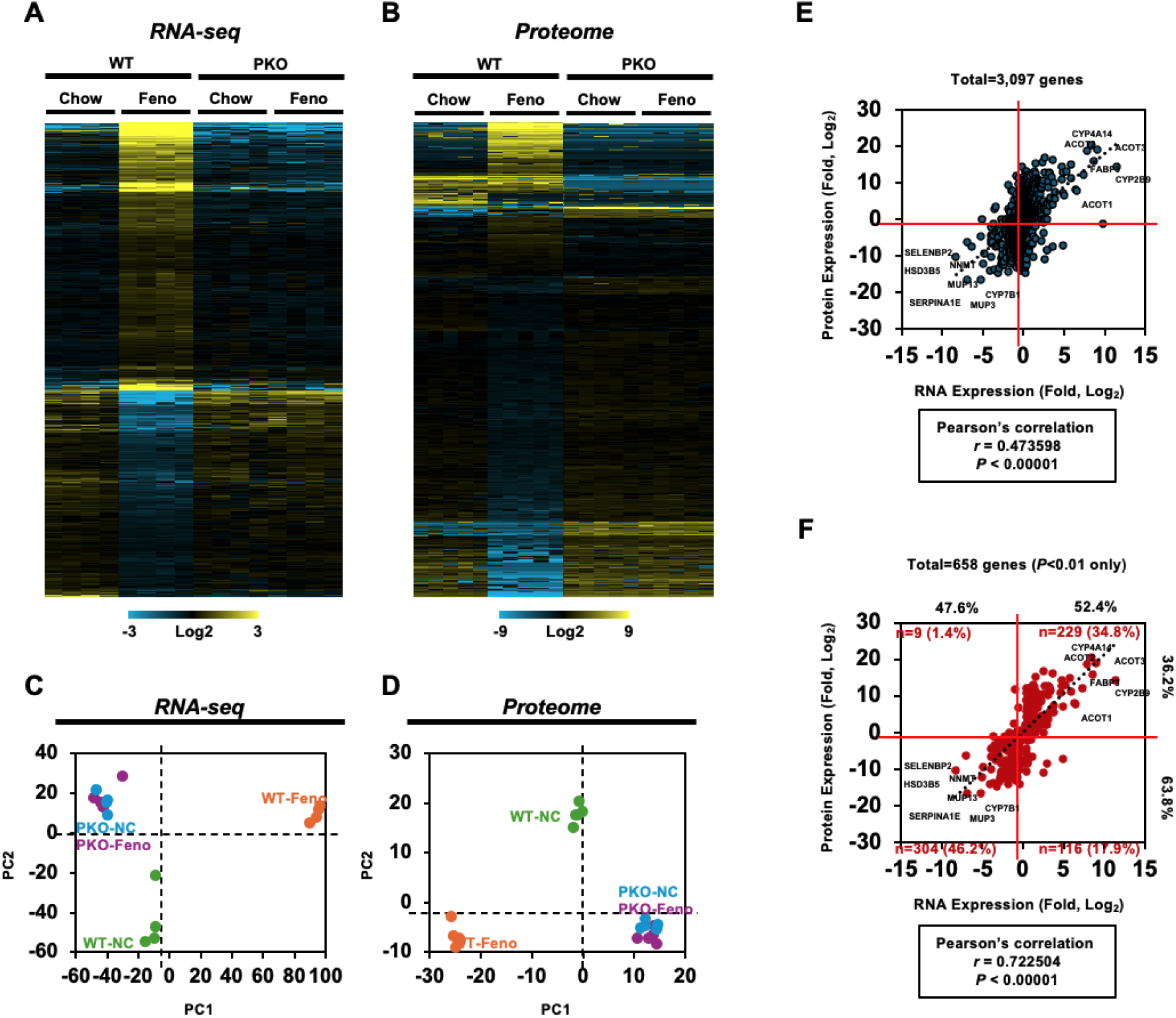
Integrative analysis of multi-omics reveals *PPARα* protein and mRNA targets. Wild-type (WT) and *PPARα*-knockout (PKO) mice were fed either a normal chow diet (Chow) or a chow diet containing 0.2% fenofibrate (Feno) for two weeks. Mice were then sacrificed for tissue collection. Liver samples were harvested and subjected to RNA sequencing and proteomic analyses (n = 5 biological replicates per group). (A, B) Hierarchical clustering heatmaps of differentially expressed genes (DEGs) across four experimental groups. Yellow indicates high expression levels, while blue represents low expression levels. (A) RNA-seq data; (B) Proteomics data. (C, D) Principal component analysis (PCA) score plots illustrating sample clustering across the four experimental groups: WT-NC (wild-type, normal chow; green), WT-Feno (wild-type, fenofibrate diet; orange), PKO-NC (*PPARα*-knockout, normal chow; blue), and PKO-Feno (*PPARα*-knockout, fenofibrate diet; purple). (C) RNA-seq data; (D) Proteomics data. (E) Integrative analysis of RNA and protein expression levels using a common gene set (total 3,097 genes). The Pearson correlation coefficient was calculated to assess the concordance between RNA-seq and proteomics data. (F) Integrative analysis of RNA and protein expression levels using a differentially expressed gene set (total 658 genes) identified as statistically significant (P < 0.01) in both RNA-seq and proteomics datasets. The Pearson correlation coefficient was calculated to assess the relationship between transcript and protein expression.

### *PPARα* Represses the Hepatic and Serum Secretome

Previous studies have provided evidence of a negative relationship between *PPARα* and secreted proteins, including coagulation factors^11,12,13,14,15,16^. Linden *et al.* (2001) reported that apolipoprotein B-containing lipoprotein secretion is increased in female *PPARα*-null mice compared to female wild-type mice, with elevated serum levels. Additionally, several studies have demonstrated that *PPARα* activation negatively regulates complement and coagulation factors, including *C2*, *C3*, *C6*, *Fga*, *F5*, and *F11*, among others^12,13,14,15,16^. Given that GO analysis identified coagulation and extracellular exosome as negative targets of *PPARα*, along with the fact that complement, coagulation factors, and exosomes are subsets of the secretome, we hypothesized that *PPARα* may regulate the overall hepatic secretome, rather than being restricted to the regulation of complement, coagulation factors, and a limited subset of secreted proteins. To validate our hypothesis, we matched the common set of 3,097 genes with the top 200 most abundant serum proteins from the Human Protein Atlas database. A total of 103 genes from this top 200 secreted protein list were identified within the common set and were subsequently used for integrative analysis (Fig. 2*A*). The Pearson correlation coefficient for these 103 genes detected in both RNA-seq and proteomic datasets revealed a moderate positive correlation (r = 0.437241) between transcript and protein expression in chow-fed and fenofibrate-fed WT livers. Among the 103 genes, 44 showed significant alterations at both the mRNA and protein levels in fenofibrate-treated WT livers. 34 out of 44 genes (77.3%), including *Serpina1e*, *Serpinf2*, *Mup8*, *C8A*, *Egfr*, *Orm1*, and others, were downregulated at both the transcript and protein levels. 4 out of 44 genes (9.1%), including *Pltp*, *Lamb3*, *Car2*, and *Actg1*, showed increased transcript levels but reduced protein levels. 4 out of 44 genes (9.1%), including *Blvrb*, *Dbi*, *Cycs*, and *H2-Q10*, were upregulated at both the transcript and protein levels. 2 out of 44 genes (4.5%), including *Pon1* and *1300017J02Rik*, showed decreased transcript levels but increased protein levels (Fig. 2*A*). Next, we further validated the mRNA levels of abundant secreted genes using qRT-PCR analysis. Consistent with our previous findings, the majority of these genes were significantly downregulated in fenofibrate-fed WT liver (Fig. 2*B*). At the protein level, most of these secreted proteins also exhibited significant downregulation (Fig. 2*C*). Since serum proteins are predominantly derived from the liver, we examined whether *PPARα* activation also affects serum protein levels (Supplemental Fig. 3). Consistent with our findings in the liver, the majority of secreted proteins in serum, including Serpins, coagulation factors, complement proteins, Apo proteins, and others, were significantly downregulated in fenofibrate-fed WT mice (Fig. 2*D*). Together, these results demonstrate that *PPARα* activation significantly downregulates the hepatic secretome at both the transcriptional and protein levels, leading to a subsequent reduction in the serum proteome. This indicates that *PPARα* broadly regulates the hepatic and serum secretome, rather than being confined to a specific subset of secreted proteins.

**Figure 2.**
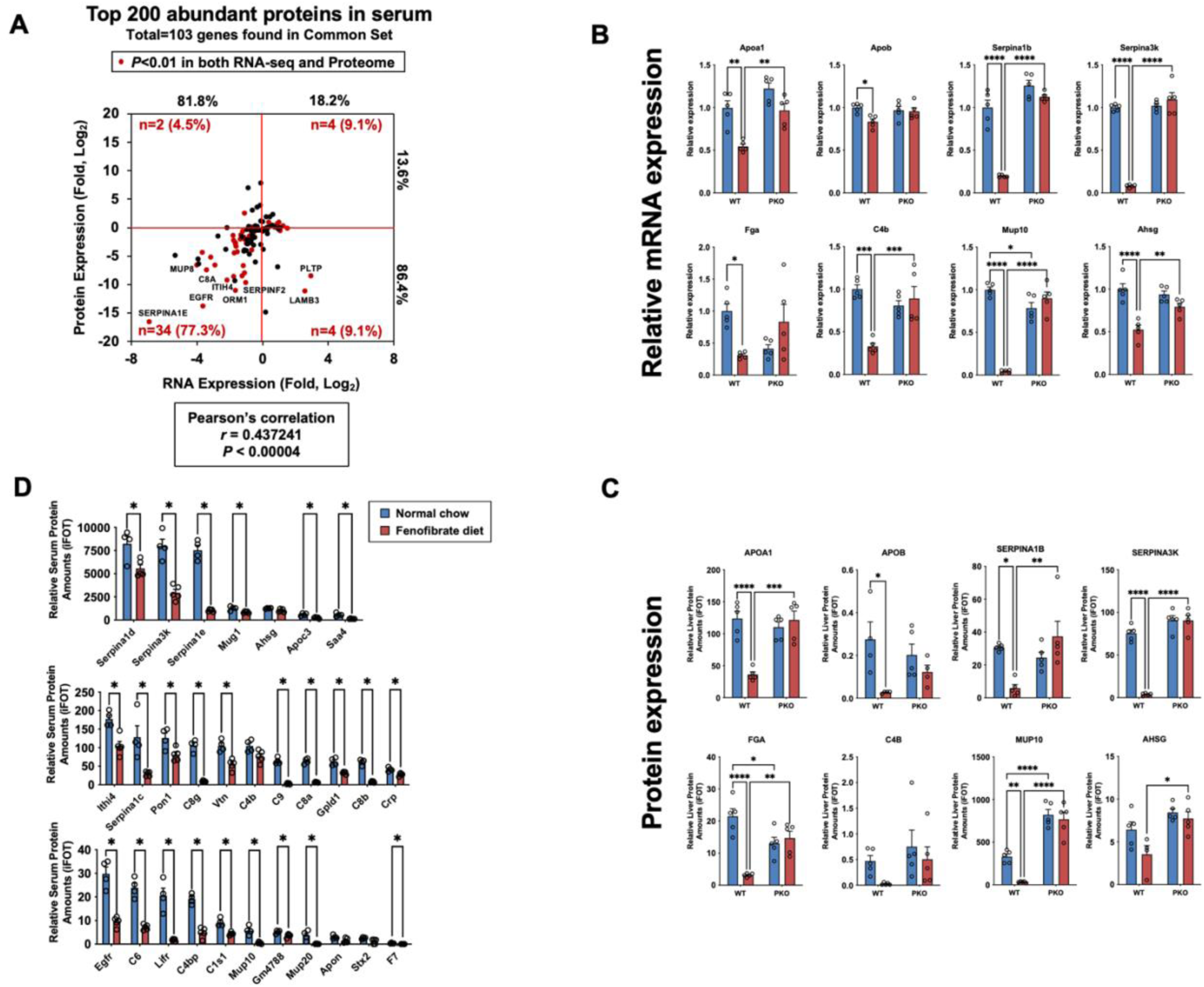
*PPARα*-Dependent Regulation of Hepatic and Circulating Secretome. WT and PKO mice were fed either a normal chow diet or a chow diet containing 0.2% fenofibrate for two weeks. Mice were then sacrificed for tissue collection. Liver and serum samples were collected for RNA sequencing, proteomic, and qPCR analyses of the liver, as well as proteomic analysis of the serum (n = 5 per group). (A) Integrative analysis of RNA and protein expression levels using the top 200 most abundant serum proteins, of which 103 genes overlapped with the common dataset. The Pearson correlation coefficient was calculated to assess the concordance between RNA-seq and proteomics data. (B) qPCR validation of mRNA expression levels for abundant secreted proteins in liver samples across the four experimental groups. (C) Relative liver protein levels (iFOT) of abundant secreted proteins across the four groups, based on proteomic analysis. (D) Serum proteomic analysis showing relative serum protein levels (iFOT) in WT-NC and WT-Feno groups. All data are presented as mean ± SEM (n = 5 per group). Statistical significance was determined using two-way ANOVA followed by Tukey’s multiple comparisons test for liver data, and multiple unpaired t-tests for serum data. *P < 0.05, **P < 0.001, ***P < 0.0001, ****P < 0.00001.

### *PPARα* Represses Endoplasmic Reticulum (ER) Protein Processing

Secreted proteins are precisely processed in the ER, where they are subjected to quality control mechanisms to ensure proper folding. Correctly folded proteins are then trafficked to the secretory pathway^23^. Since we confirmed that secretome genes are downregulated in fenofibrate-fed liver, we investigated whether ER protein processing genes are also regulated by *PPARα* activation. First, we filtered and identified 89 ER protein processing-related genes within the common set, and these genes were used for integrative analysis (Fig. 3*A*). The Pearson correlation coefficient for these 89 genes detected in both RNA-seq and proteomic datasets revealed a moderate positive correlation (r = 0.509431) between transcript and protein expression in chow-fed and fenofibrate-fed WT livers. Among the 89 genes, 33 genes exhibited significant changes at both the transcript and protein levels in fenofibrate-fed WT livers: 30 out of 33 genes (90.9%), including *Ssr3*, *Sec24d*, *Tram1*, *Ero1lb*, *Hyou1*, and others, were downregulated at both the transcript and protein levels. 2 out of 33 genes (6.06%), including *Dnaja2* and *Nsfl1c*, were upregulated at both the transcript and protein levels. 1 out of 33 genes (3%), *Bax*, exhibited increased transcript levels but decreased protein levels (Fig. 3*A*). ER performs multiple functions depending on its structural organization. For example, rough ER sheet structures decorated by ribosomes are specialized for enhanced protein synthesis and secretion, whereas smooth ER tubular structures are predominantly associated with lipid and steroid synthesis^24,25,26,27,28,29^. To assess potential *PPARα*-induced changes in ER morphology, which may reflect shifts in its functional specialization, we performed transmission electron microscopy (TEM) of liver sections of chow-fed and fenofibrate-fed WT mice. As suggested by the origin of its name, *PPARα* (Peroxisome Proliferator-Activated Receptor Alpha), a notable increase in peroxisomes and mitochondria was observed in fenofibrate-fed hepatocytes compared to those in chow-fed hepatocytes (Fig. 3*B*)^30^. Interestingly, hepatocytes from fenofibrate-fed mice exhibited a significant reduction in parallel organized rough ER sheet structures compared to chow-fed hepatocytes. Instead, the rough ER is organized as single sheet surround the peroxisomes and mitochondria. We also detected higher levels of tubular ER. These morphological changes are consistent with reduced rough ER dependent protein processing capacity. Additionally, the rough ER enveloped mitochondria and peroxisomes, have been suggested to regulate lipid oxidation^31^. Therefore, the induction of these structures by *PPARα* activation is consistent with the role of *PPARα* in lipid regulation (Fig. 3*B*). Next, we further validated the mRNA levels of ER protein processing genes in the liver using qPCR analysis. Consistent with our previous findings, the majority of these genes, including those involved in translocon function (*Sec61a1*, *Sec63*, *Ssr1*), OST complex activity (*Rpn1*, *Ddost*), glycosylation (*Uggt1*), glycoprotein folding (*Calr*, *Pdia3*), ER luminal chaperone activity (*Hspa5*, *Hsp90b1*), lectin function (*Lman1*, *Lman2*), vesicle trafficking (*Sec24d*), and oxidoreduction (*Ero1lb*, *Pdia4*, *Pdia6*), were significantly downregulated in fenofibrate-fed WT liver (Fig. 3*C*). At the protein level, most of these ER processing proteins also exhibited significant downregulation (Fig. 3*D*). These results demonstrate that *PPARα* activation reduces the transcription and translation of liver protein processing-related mRNAs, coinciding with significant structural changes in the ER.

**Figure 3.**
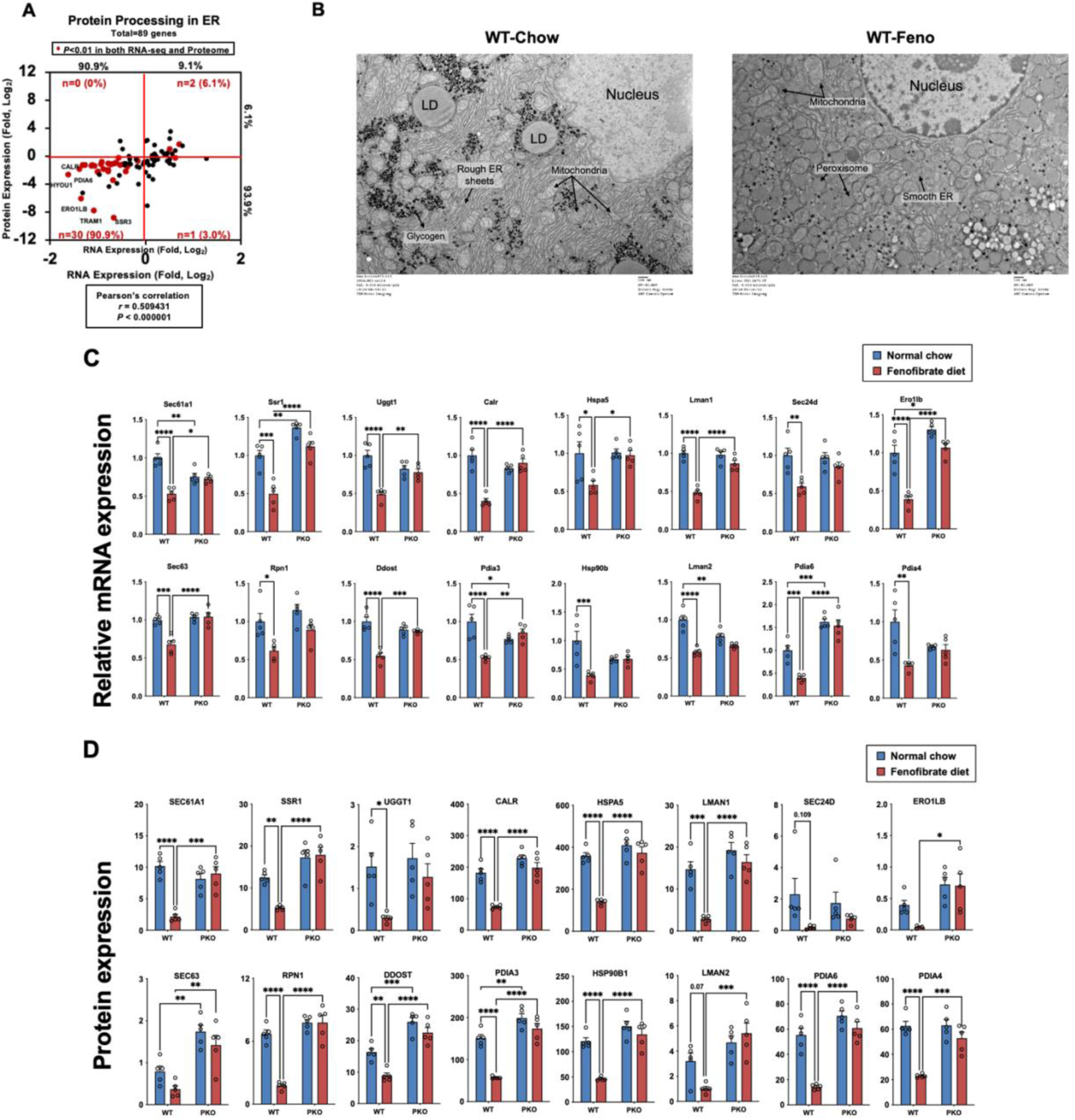
*PPARα* Modulates Endoplasmic Reticulum Architecture and Functions. WT and PKO mice were fed either a normal chow diet or a chow diet containing 0.2% fenofibrate for two weeks. Mice were then sacrificed for tissue collection. Liver samples were harvested and subjected to RNA sequencing, proteomic, and qPCR analyses (n = 5 per group). For EM images, livers were perfused via the portal vein with saline, followed by fixation. The liver tissues were then sectioned into 300-μm slices and further incubated in a secondary fixative. For TEM imaging, samples were embedded in resin, sectioned into ultrathin slices and examined using transmission electron microscope. (A) Integrative analysis of RNA and protein expression levels using a protein processing–related gene set, with 89 genes overlapping the common dataset. The Pearson correlation coefficient was calculated to assess the concordance between RNA-seq and proteomics data. (B) Transmission electron microscopy (TEM) images of liver tissue from WT-Chow (left) and WT-Feno (right) mice. Scale bar: 500 nm. (C) qPCR validation of mRNA expression levels for protein processing-related gene sets in liver samples across the four experimental groups. (D) Relative liver protein levels (iFOT) of protein processing-related gene sets across the four groups, based on proteomic analysis. These include translocon components (*Sec61a1*, *Sec63*, *Ssr1*), the ribosome-binding gene *Rrbp1*, OST complex members (*Stt3a*, *Rpn1*, *Ddost*), the glycosylation gene *Uggt1*, glycoprotein folding genes (*Calr*, *Pdia3*), ER luminal chaperones (*Hspa5*, *Hsp90b1*), lectins (*Lman1*, *Lman2*), the vesicle trafficking gene *Sec24d*, and oxidoreduction-related genes (*Ero1lb*, *Pdia4*, *Pdia6*). All data are presented as mean ± SEM (n = 5 per group). Statistical significance was determined using two-way ANOVA followed by Tukey’s multiple comparisons test for liver data. *P < 0.05, **P < 0.001, ***P < 0.0001, ****P < 0.00001.

### *PPARα* Enhances Protein Degradation to Facilitate Amino Acid Recycling

The proteostasis network consists of protein processing, secretion, degradation, and synthesis, maintaining protein homeostasis and cellular function^32^. In our previous study, we demonstrated that hepatic autophagy is directly regulated by the nuclear receptors *PPARα* and *FXR*^17^. In eukaryotic cells, old, damaged, or misfolded proteins are degraded through either the ubiquitin– proteasome system or the autophagic–lysosomal pathway to support amino acid recycling under nutrient-deprived conditions^32,33^. Given that *PPARα* promotes autophagy-related degradation, we next investigated the proteasome activity, the other axis of protein degradation, in *PPARα*-activated liver. To evaluate the effect of *PPARα* activation on proteasome function, we measured the proteolytic degradation activity of polyubiquitinated substrates in liver samples from each experimental group using a proteasome activity assay kit (UBPbio, Cat# J4110). Fenofibrate-fed WT livers exhibited higher chymotrypsin-like activity of the 26S proteasome compared to other groups, indicating that *PPARα* activation enhances proteasome function (Fig. 4*A*). As the proteasome consists of a 20S core and 26S regulatory components—where the 20S core is formed by PSMA (two outer α-rings) and PSMB (two inner β-rings) families, and the 26S regulator includes PSMC (ATP-dependent hexameric ATPase ring) and PSMD (ATP-independent lid) families—we assessed the mRNA expression levels of representative subunits from each family. The expression levels of PSMA, PSMB, and PSMC subunits were generally comparable between chow-fed and fenofibrate-fed WT groups. In contrast, several PSMD subunits—including *Psmd1*, *Psmd3*, *Psmd4*, *Psmd5*, *Psmd8*, *Psmd10*, *Psmd11*, *Psmd13*, and *Psmd*14—exhibited statistically significant increases in mRNA expression in the fenofibrate-fed WT group (Fig 4*B* and *C*). These findings indicate that *PPARα* activation enhances proteasome activity, likely through the upregulation of PSMD family members, which constitute the ATP-independent lid of the 26S proteasome.

**Figure 4.**
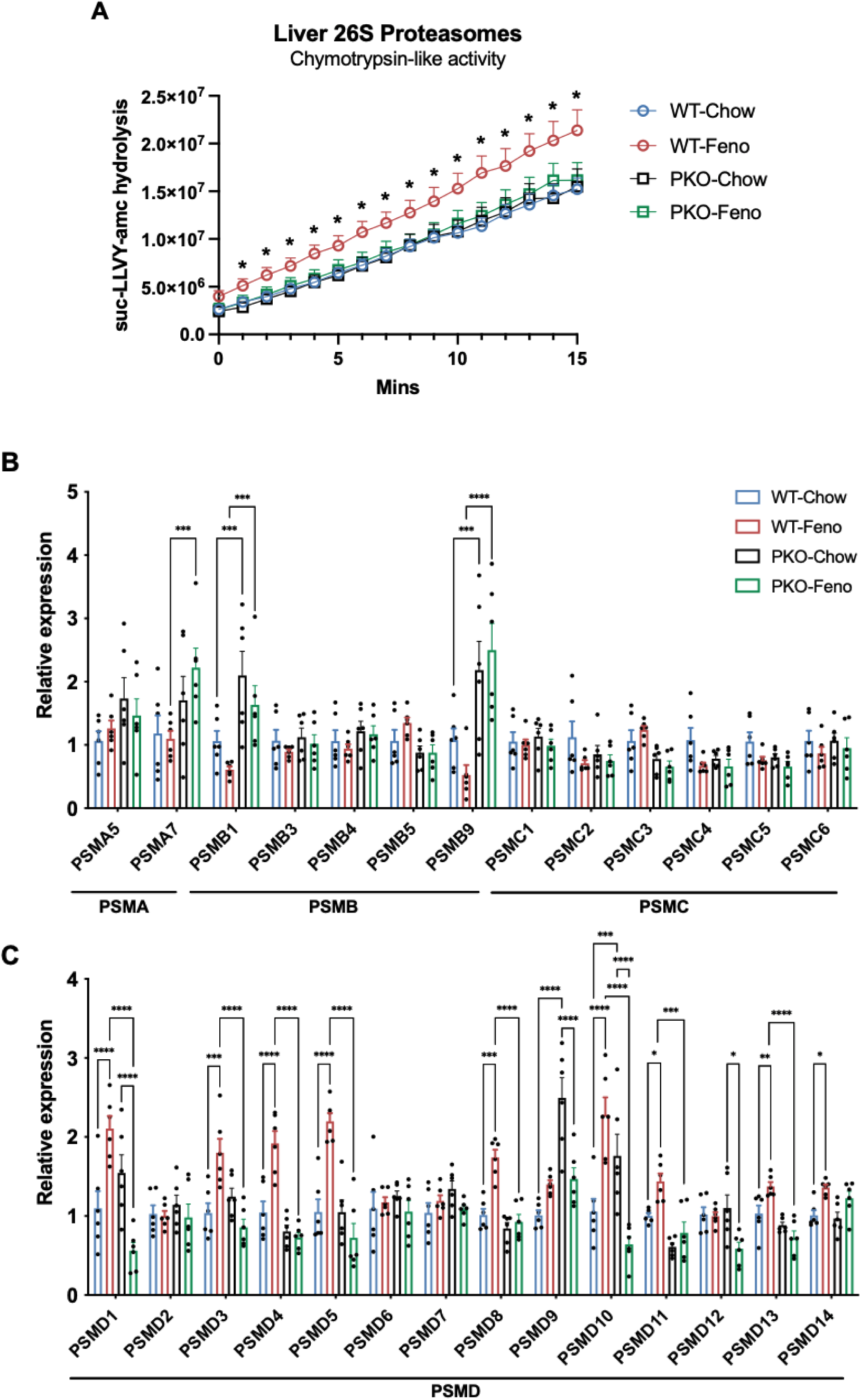
*PPARα* Stimulates Proteasomal Protein Degradation. WT and PKO mice were fed either a normal chow diet or a chow diet containing 0.2% fenofibrate for two weeks. Mice were then sacrificed for tissue collection. Liver samples were harvested and 50mg of tissues were homogenized in RIPA buffer for measurement of chymotrypsin-like activity of the 26S proteasome. A separate portion of the tissue was homogenized in TRI Reagent for RNA isolation, followed by qPCR analysis. (A) Chymotrypsin-like activity of hepatic 26S proteasomes measured across the four experimental groups. (B) qPCR analysis of mRNA expression levels for proteasome subunit families (PSMA, PSMB, PSMC) in liver samples from the four experimental groups. (C) qPCR analysis of mRNA expression levels for proteasome subunit PSMD family in liver samples from the four experimental groups. All data are presented as mean ± SEM (n = 6 per group). Statistical significance was determined using two-way ANOVA followed by Tukey’s multiple comparisons test for liver data, and multiple unpaired t-tests for liver chymotrypsin-like activity data. *P < 0.05, **P < 0.001, ***P < 0.0001, ****P < 0.00001.

### *PPARα* controls protein synthesis or translation initiation

Protein synthesis is a fundamental process required for maintaining proteostasis. Based on the integrative analysis of 3,097 genes shared between RNA-seq and proteomics datasets, we identified genes whose RNA expression remained largely unchanged, while their protein expression exhibited significant changes, suggesting the involvement of post-transcriptional regulation, particularly in the efficiency of protein translation (Fig. 1*E*). In addition, our integrative analysis of ribosomal proteins provided a clear example of post-transcriptional regulation. Among the 62 ribosomal proteins identified in the common set, 37 ribosomal proteins exhibited significant changes at both the transcript and protein levels in fenofibrate-fed WT livers. Interestingly, all 37 ribosomal proteins showed upregulated RNA expression but significantly downregulated protein expression, indicating post-transcriptional regulation that may directly impact protein synthesis rates (Supplemental Fig. 4). To further investigate protein translation efficiency, we conducted assays to measure *de novo* protein synthesis rates. To assess short-term protein synthesis, we performed a SUnSET assay following a well-established protocol (Supplemental Fig. 5*A*)^34^. Fenofibrate-fed WT liver exhibited a general reduction in puromycin-labeled proteins compared to chow-fed WT liver, indicating a lowered short-term protein synthesis rate upon *PPARα* activation. In contrast, this reduction was not observed in PKO mice, as puromycin incorporation remained comparable between fenofibrate- and chow-fed PKO livers (Fig. 5*A*). To assess long-term protein synthesis, we performed a deuterium (D_2_O) labeling assay in mice for 72 hours (Supplemental Fig. 5*B*). The average liver fractional synthesis rate in fenofibrate-fed WT mice was significantly reduced compared to chow-fed WT mice, whereas this effect was not observed in PKO mice, indicating that *PPARα* activation lowers long-term protein synthesis rates, leading to a slower protein turnover in the liver (Fig. 5*B*). We next examined whether the reduction in protein synthesis is associated with alterations in the translational machinery, including translation initiation and elongation factors. eIF2A phosphorylation (Ser51) exhibited an approximately 3.5-fold increase in fenofibrate-fed WT livers compared to the chow-fed condition, whereas this effect was not observed in PKO mice. In contrast, eIF4E phosphorylation (Ser209), another rate-limiting factor in translation initiation, was reduced by approximately 30% under fenofibrate-fed conditions compared to chow-fed livers. The phosphorylation of eEF2 (Thr56), a key regulator of translation elongation, remained unchanged between chow- and fenofibrate-fed WT livers. Similarly, phosphorylation of RPS6 (Ser235/236) showed no significant difference between the two conditions (Fig. 5*C* and *D*). Collectively, these findings demonstrate that *PPARα* activation reduces global protein translation rates in both short-term and long-term protein synthesis, likely by inhibiting translational initiation rather than elongation steps.

**Figure 5.**
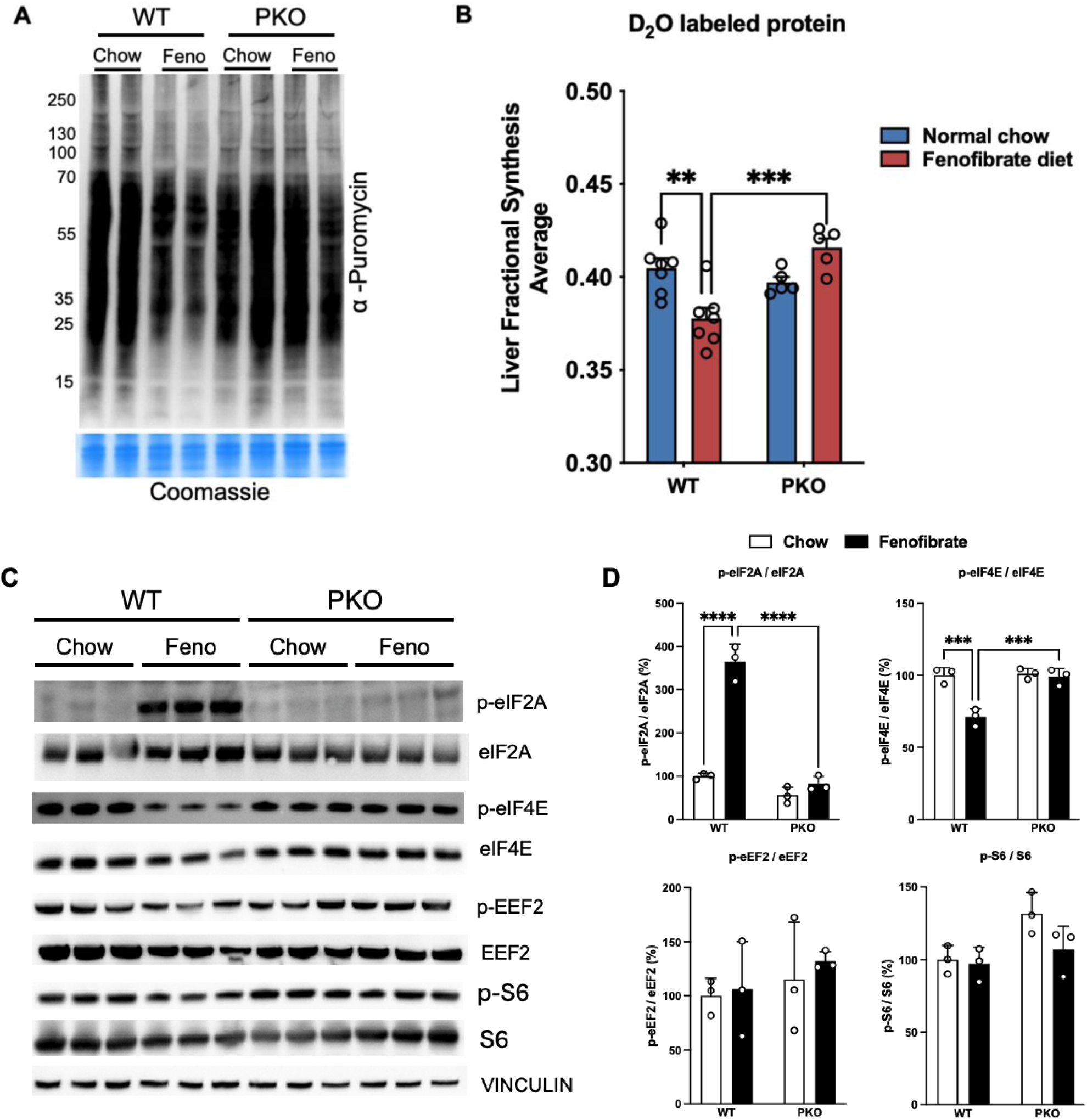
*PPARα* Represses Global Protein Translation Through eIF2α Phosphorylation. WT and PKO mice were fed either a normal chow diet or a chow diet containing 0.2% fenofibrate for two weeks. To assess short-turnover protein synthesis rates using the SUnSET assay, animals were intraperitoneally injected with puromycin at a dose of 40 nmol/g body weight. After 45 minutes, mice were sacrificed, and livers were harvested. To assess long-turnover protein synthesis rates using the heavy water (D₂O) labeling assay, mice received a priming intraperitoneal (IP) injection of 100% D₂O, followed by continuous access to drinking water containing 8% D₂O for three days. After the labeling period, mice were sacrificed, and liver tissues were harvested and processed for liquid chromatography–mass spectrometry (LC-MS) analysis. (A) Western blot–based SUnSET analysis of liver samples to assess short-turnover protein synthesis across the four experimental groups. Coomassie staining was used as a loading control (n = 2 per group). (B) Heavy water (D₂O) labeling analysis of long-turnover protein synthesis in the liver: average fractional synthesis rates of total hepatic proteins across the four groups (n = 7 for WT, n = 5 for PKO). (C) Western blot analysis of translational machinery proteins in liver samples from the four experimental groups (n = 3 per group), including phosphorylated and total forms of eIF2A, eIF4E, EEF2, and S6. Vinculin was used as a loading control. (D) Quantification of each translational machinery protein and relative expression levels of phosphorylated proteins normalized to their corresponding total protein levels. All data are presented as mean ± SEM (n = 3 per group). Statistical significance was determined using two-way ANOVA followed by Tukey’s multiple comparisons test for liver data. *P < 0.05, **P < 0.001, ***P < 0.0001, ****P < 0.00001.

### *PPARα* phosphorylates eIF2A via the iron – HRI axis

Phosphorylation of eIF2A is known to repress general translation^35^. Four upstream kinases— General control nonderepressible 2 kinase (GCN2), PKR-like endoplasmic reticulum kinase (PERK), Protein kinase R (PKR), and Heme-regulated inhibitor kinase (HRI)—have been identified as regulators of eIF2A phosphorylation^36^. Given the significant increase in eIF2A phosphorylation observed in fenofibrate-fed WT liver, we assessed the activity of these four kinases by measuring their phosphorylated form relative to total protein levels (Fig. 6*A*). Phosphorylation levels of GCN2 remained unchanged across all groups, while PERK and PKR phosphorylation was significantly reduced in fenofibrate-fed WT livers. In contrast, HRI phosphorylation was markedly increased in this group, suggesting that HRI activation mediates the elevated eIF2α phosphorylation observed in fenofibrate-fed WT livers (Fig. 6A and B). As heme deficiency leads to HRI phosphorylation, we next examined hepatic heme levels in each group^37^. Heme levels were reduced in fenofibrate-fed WT livers compared to chow-fed WT livers, although the difference approached but did not reach statistical significance (P = 0.058). No such change was observed in PKO mice (Fig. 6C, left). To investigate whether the reduction in heme levels was due to iron availability, we measured iron levels in both liver and serum. Interestingly, hepatic iron levels were significantly decreased in fenofibrate-fed WT mice compared to chow-fed WT mice, whereas serum iron levels remained unchanged between the two groups (Fig. 6*C*, middle and right). To further explore the molecular basis of this iron depletion, we examined the expression of iron metabolism-related genes. The mRNA expression of transferrin (*Tf*) and transferrin receptor 1 (*Tfrc*), was dramatically downregulated in fenofibrate-fed WT liver (Fig. 6D), whereas this change was not observed in PKO mice. In contrast, ferritin heavy polypeptide 1 (*Fth1*) mRNA expression was significantly increased in fenofibrate-fed WT liver compared to chow-fed WT liver whereas ferritin light polypeptide 1 (*Ftl1*) mRNA expression was comparable between chow-fed and fenofibrate-fed WT livers. The expression levels of *hepcidin*, ceruloplasmin (*Cp*), ferroportin 1 (*Fpn1*), and divalent metal transporter 1 (*Dmt1*) remained comparable between fenofibrate- and chow-fed WT livers (Fig. 6*D*). Together, our findings indicate that *PPARα* activation reduces hepatic heme and iron levels, leading to increased HRI phosphorylation, which may be attributed to the downregulation of *Tf* and *Tfrc* expression in the liver.

**Figure 6.**
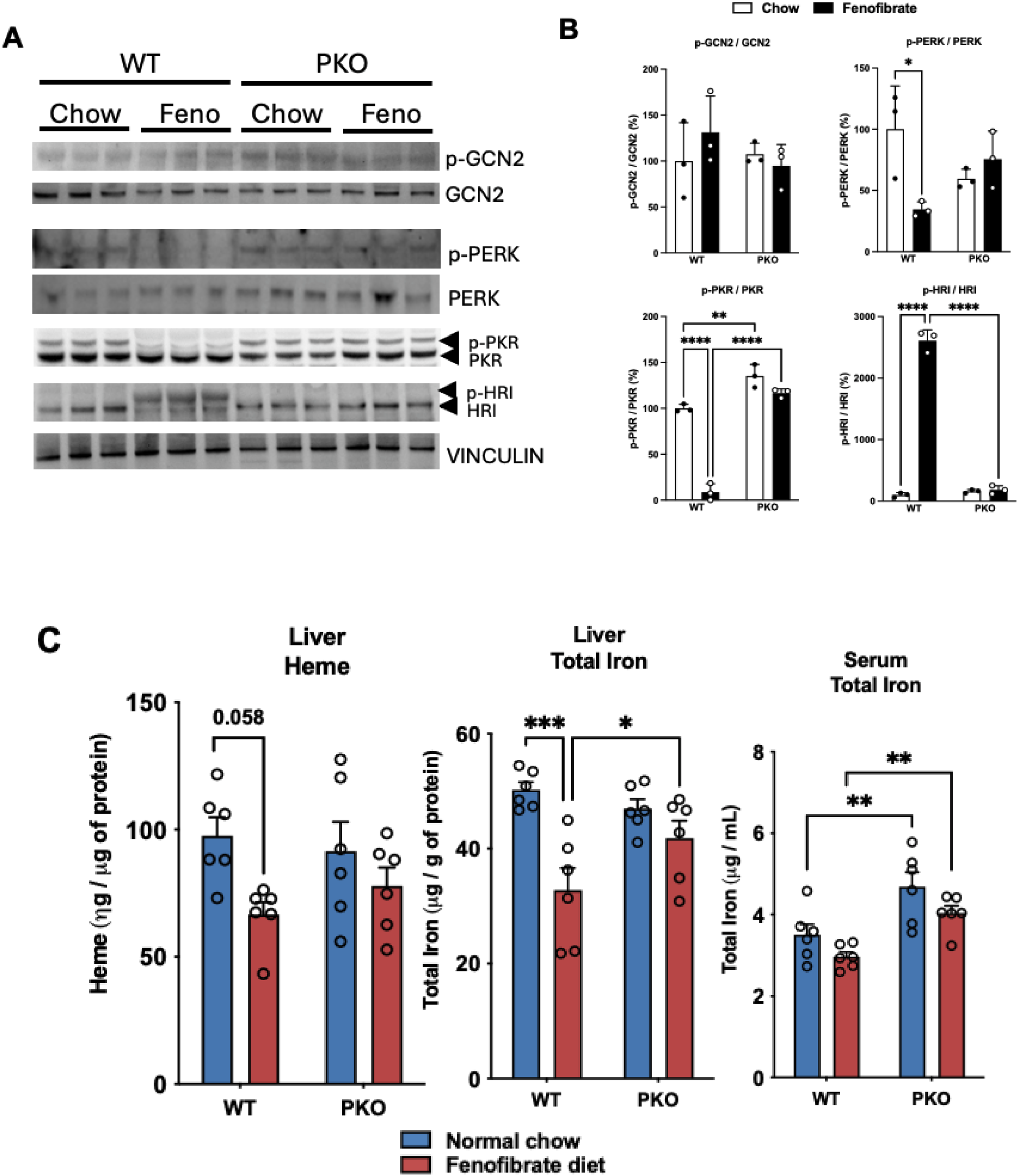
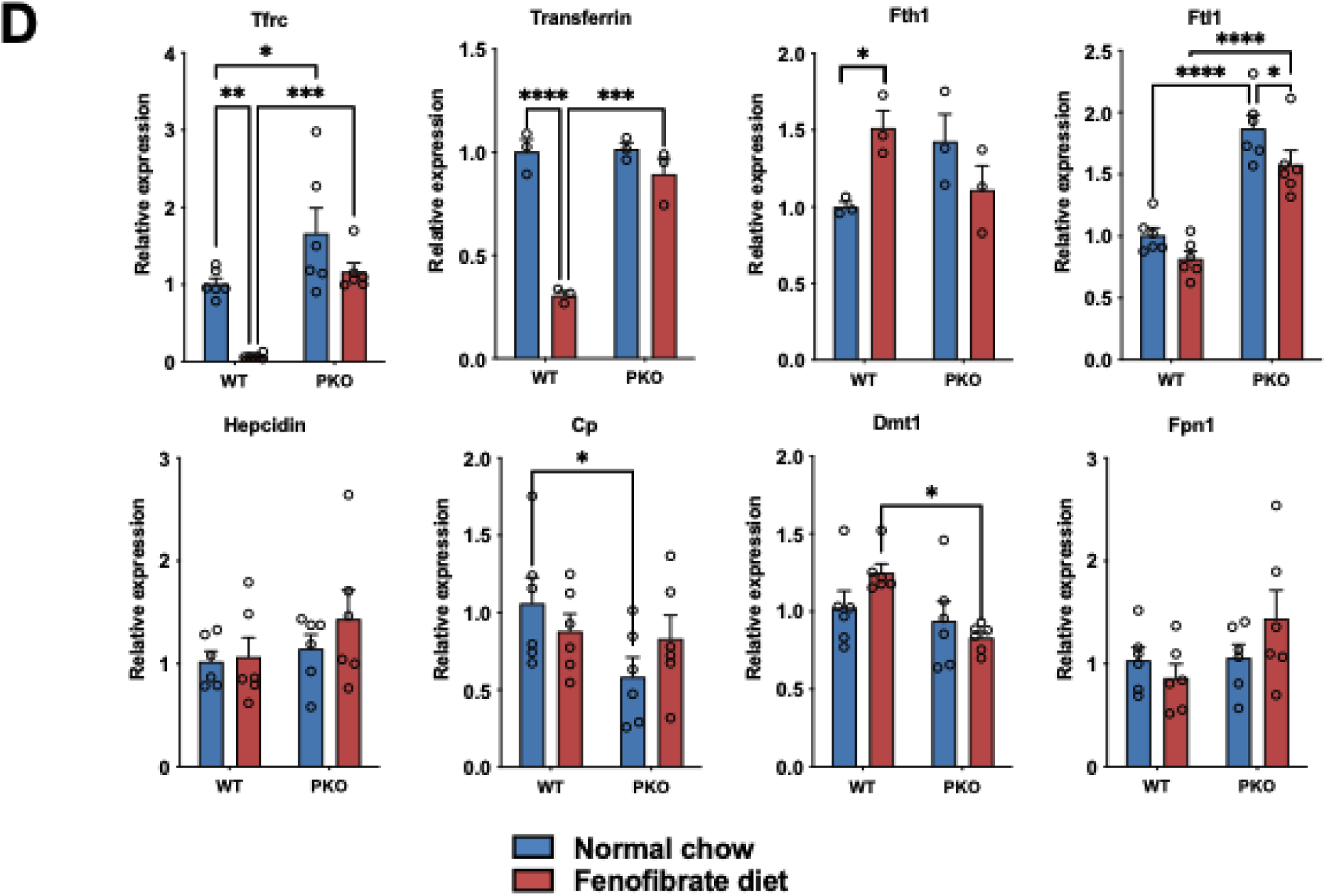
*PPARα* Activates the Iron–HRI–eIF2α Signaling Pathway. WT and PKO mice were fed either a normal chow diet or a chow diet containing 0.2% fenofibrate for two weeks. Mice were then sacrificed for tissue collection. Liver samples were harvested and 50mg of tissues were homogenized in RIPA buffer for measurement of heme and total iron contents as well as western blot analysis. A separate portion of the tissue was homogenized in TRI Reagent for RNA isolation, followed by qPCR analysis. (A) Western blot analysis of upstream kinase proteins of eIF2A in liver samples from the four experimental groups (n = 3 per group), including phosphorylated and total forms of GCN2, PERK, PKR, and HRI. Vinculin was used as a loading control. (B) Quantification of each translational machinery protein and relative expression levels of phosphorylated proteins normalized to their corresponding total protein levels. (C) Measurement of liver heme content (left), total liver iron content (middle), and serum iron content (right) across the four experimental groups (n = 6 per group). (D) qPCR analysis of mRNA expression levels for iron metabolism-related genes in liver samples from the four experimental groups (n = 6 per group). All data are presented as mean ± SEM. Statistical significance was determined using two-way ANOVA followed by Tukey’s multiple comparisons test for liver data. *P < 0.05, **P < 0.001, ***P < 0.0001, ****P < 0.00001.

## Discussion

*PPARα* is a well-characterized nuclear receptor that is activated in the fasting state and plays a central role in regulating peroxisomal and mitochondrial fatty acid oxidation, ketogenesis, gluconeogenesis, and other metabolic pathways. These functions highlight the critical role of *PPARα* in maintaining glucose and lipid homeostasis^4,6,8,38^. While extensive studies have documented its role in glucose and lipid metabolism, the involvement of *PPARα* in protein metabolism remains less explored. However, existing evidence suggests that *PPARα* directly or indirectly regulates genes involved in proteostasis, including those associated with amino acid metabolism, secreted proteins and autophagy-mediated protein clearance^11,12,13,14,15,16,32^. In this study, we uncovered a previously unrecognized role of *PPARα* in maintaining protein homeostasis.

We observed that chronic pharmacological activation of *PPARα* induced extensive changes in the hepatic transcriptome and proteome. As expected, upregulated genes were enriched in pathways related to mitochondrial and peroxisomal fatty acid metabolism. In contrast, downregulated genes were associated with pathways involved in protein secretion, ER protein processing, complement and coagulation cascades, and iron binding. Though it is well-established that *PPARα* negatively regulates certain complement and coagulation-related genes, our findings extend this understanding by demonstrating that *PPARα* broadly suppresses the secretome in liver and circulation. Previous results suggested that transcriptional regulation of coagulation factor genes may involve direct competition at the consensus IR-1 FXR response element^16^. *While the precise mechanisms underlying the regulation of secretome genes require further investigation, the findings from this study suggest that one potential mechanism could be involved in modulation of secreted protein processing within ER.* The structural shift of the ER from parallel sheets to tubular forms in fenofibrate-fed livers supports a reduction in protein processing capacity, while indicating enhanced lipid metabolism and fatty acid oxidation. This is also supported by the increased number of lipid droplets and the presence of tubular ER structures closely associated with mitochondria and peroxisomes. From a proteostasis perspective, protein secretion accounts for the highest gene expression priority in the liver, with approximately 40% of hepatic mRNAs encoding secreted proteins. Given that the liver contributes an estimated 2–30% of total body energy expenditure in humans, hepatic secretion represents a highly energy-demanding process^39,40^. Based on our findings, hepatic secretion is reciprocally regulated by *PPARα* through direct transcriptional control of secreted genes, modulation of ER protein maturation processes, and translational regulation—altogether supporting its central role in energy conservation.

Protein clearance is primarily mediated by two independent pathways: the autophagy–lysosome system and the ubiquitin–proteasome system. While the autophagy pathway is rapidly activated during nutrient starvation and is responsible for degrading long-lived proteins, aggregated proteins, and intracellular organelles, the ubiquitin–proteasome system predominantly degrades short-lived, misfolded, and damaged proteins^41,42,43,44^. Previously, we identified that pharmacological activation of *PPARα* can directly promote autophagy by binding to the direct repeat 1 (DR1) motif on the promoters of key autophagy-related genes, presumably facilitating the degradation of long-lived proteins, aggregated proteins, and damaged intracellular organelles^17^. Our findings show that *PPARα* activation enhances proteasome activity, likely through upregulation of the PSMD family, suggesting that *PPARα* contributes to the degradation of short-lived, misfolded, and damaged proteins as well, and facilitates amino acid recycling through the coordinated activation of both autophagy and proteasome pathways. It would be interesting to investigate which types of proteins are preferentially targeted by these degradation systems.

The mammalian target of rapamycin (mTOR) is a well-established master regulator of protein synthesis, and its activity is modulated by a variety of signals, including hormones, growth factors, and nutrient availability^45^. Interestingly, our findings demonstrate that the dramatic reduction in translation observed upon pharmacological activation of *PPARα* is not mediated through the mTOR pathway but rather involves a direct effect on translation initiation factors. The increase in HRI phosphorylation induced by iron reduction upon *PPARα* activation was entirely unexpected. However, our findings are supported by previous reports showing that *PPARα* mitigates iron overload-induced ferroptosis by reducing hepatic iron content, likely through the downregulation of transferrin expression^46^. This consistency supports the notion that *PPARα* plays a role in iron homeostasis, which may, in turn, promote HRI activation under specific physiological conditions, particularly those related to nutrient deprivation. The observation of reduced hepatic iron levels— but not serum iron—in fenofibrate-fed WT mice, along with the downregulation of transferrin and transferrin receptor 1 further supports this notion and provides additional evidence for the role of *PPARα* in regulating iron metabolism in hepatocytes. The detailed molecular mechanisms underlying this regulation remain to be investigated.

In conclusion, our study highlights novel functions of *PPARα* in hepatic proteostasis, demonstrating its role in conserving protein energy balance by restricting unnecessary protein loss through synthesis and secretion, while enhancing amino acid recycling via coordinated activation of autophagy and proteasome-mediated degradation pathways.

---

**Supplemental Figure 1.**
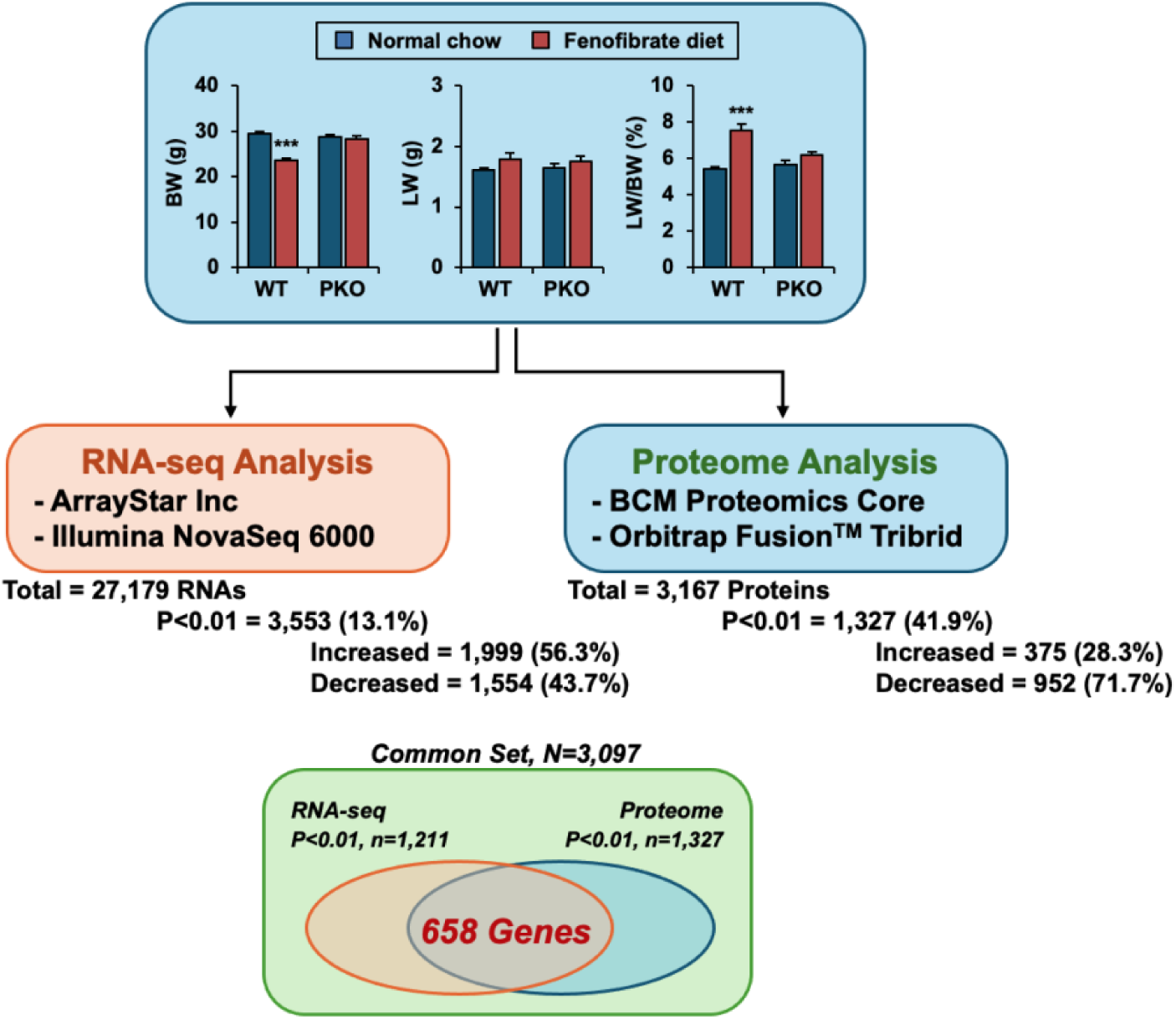
Integrative Transcriptomic and Proteomic Analyses Reveal Common Fenofibrate-Responsive Genes. Overview of integrative transcriptomic and proteomic analyses in WT and PKO mice under chow or fenofibrate diet conditions. Body weight (BW), liver weight (LW), and liver-to-body weight ratio (LW/BW) were measured across wild-type (WT) and *PPARα*-knockout (PKO) mice fed either a normal chow or 0.2% fenofibrate-containing diet. Liver samples were subjected to RNA-seq (left) and proteomic (right) analyses. RNA-seq was performed using the Illumina NovaSeq 6000 platform, identifying a total of 27,179 RNAs. Differential expression analysis (P < 0.01) revealed 3,553 genes (1,999 upregulated and 1,554 downregulated). Proteomic analysis was conducted using Orbitrap Fusion™ Tribrid mass spectrometry at the BCM Proteomics Core, quantifying 3,167 proteins. Of these, 1,327 proteins were significantly changed (P < 0.01), with 375 upregulated and 952 downregulated. A total of 3,097 genes were detected in both datasets, with 658 genes significantly differentially expressed in both RNA-seq and proteomics datasets.

**Supplemental Figure 2.**
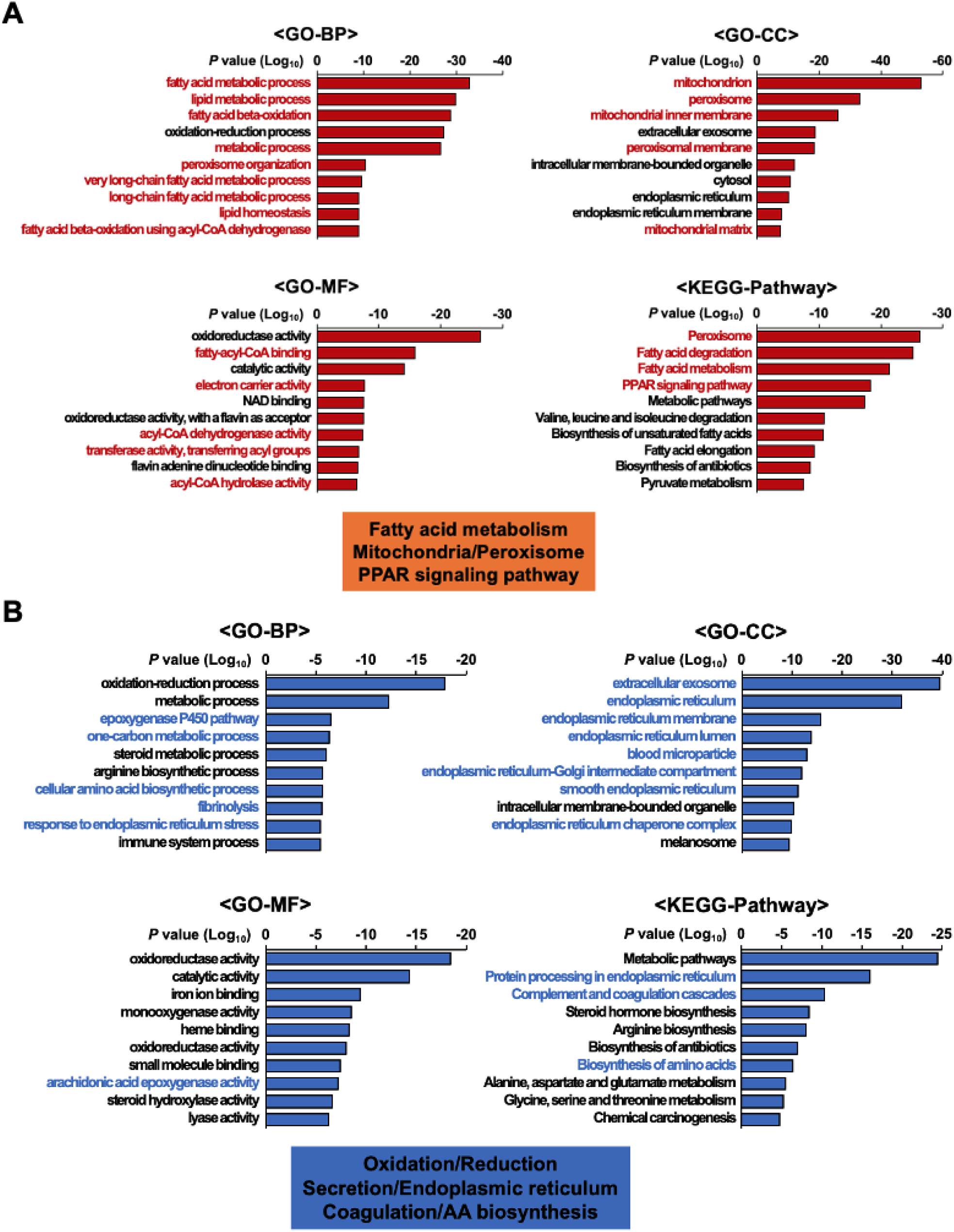
Gene Ontology (GO) analysis of 658 genes significantly differentially expressed in both RNA-seq and proteomics datasets. (A) GO enrichment analysis of upregulated genes revealed top biological processes (BP), cellular components (CC), molecular functions (MF), and KEGG pathways associated with fatty acid metabolism, mitochondrial and peroxisomal function, and PPAR signaling. (B) GO enrichment analysis of downregulated genes identified key categories related to oxidation-reduction processes, secretion and endoplasmic reticulum function, coagulation, and amino acid biosynthesis.

**Supplemental Figure 3.**
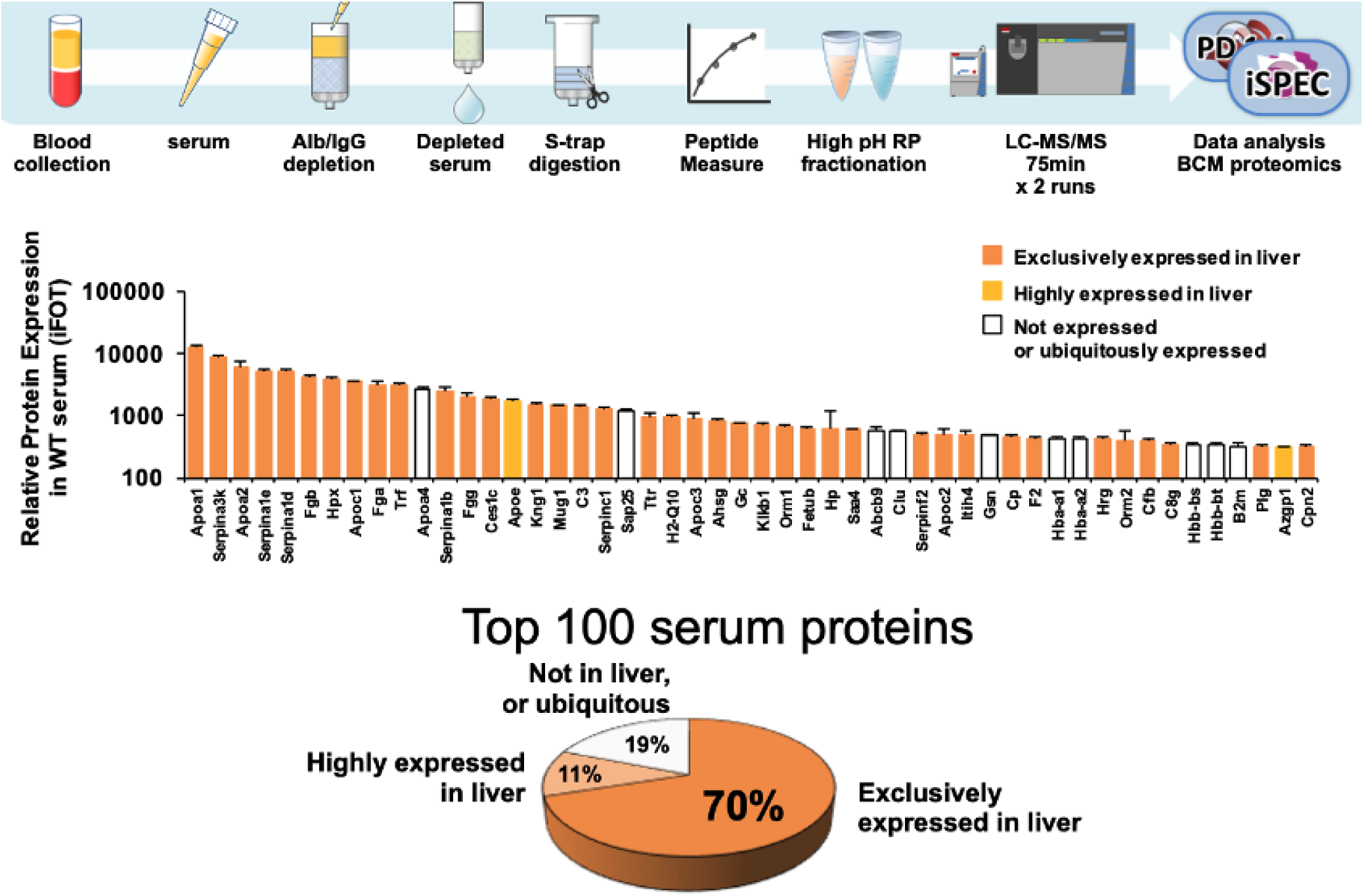
Liver as the Primary Source of the Top 100 Circulating Serum Proteins. Schematic of the serum proteomics workflow. Following blood collection, serum was subjected to albumin/IgG depletion, S-Trap digestion, peptide measurement, and high-pH reverse-phase (RP) fractionation. Peptides were analyzed using LC-MS/MS (75 min × 2 runs), and data were processed via BCM Proteomics Core using iSPEC. Bar graph showing relative expression levels of the top 100 most abundant serum proteins. Proteins are color-coded by expression profile: exclusively expressed in liver (orange), highly expressed in liver (yellow), and not expressed or ubiquitously expressed (white).

**Supplemental Figure 4.**
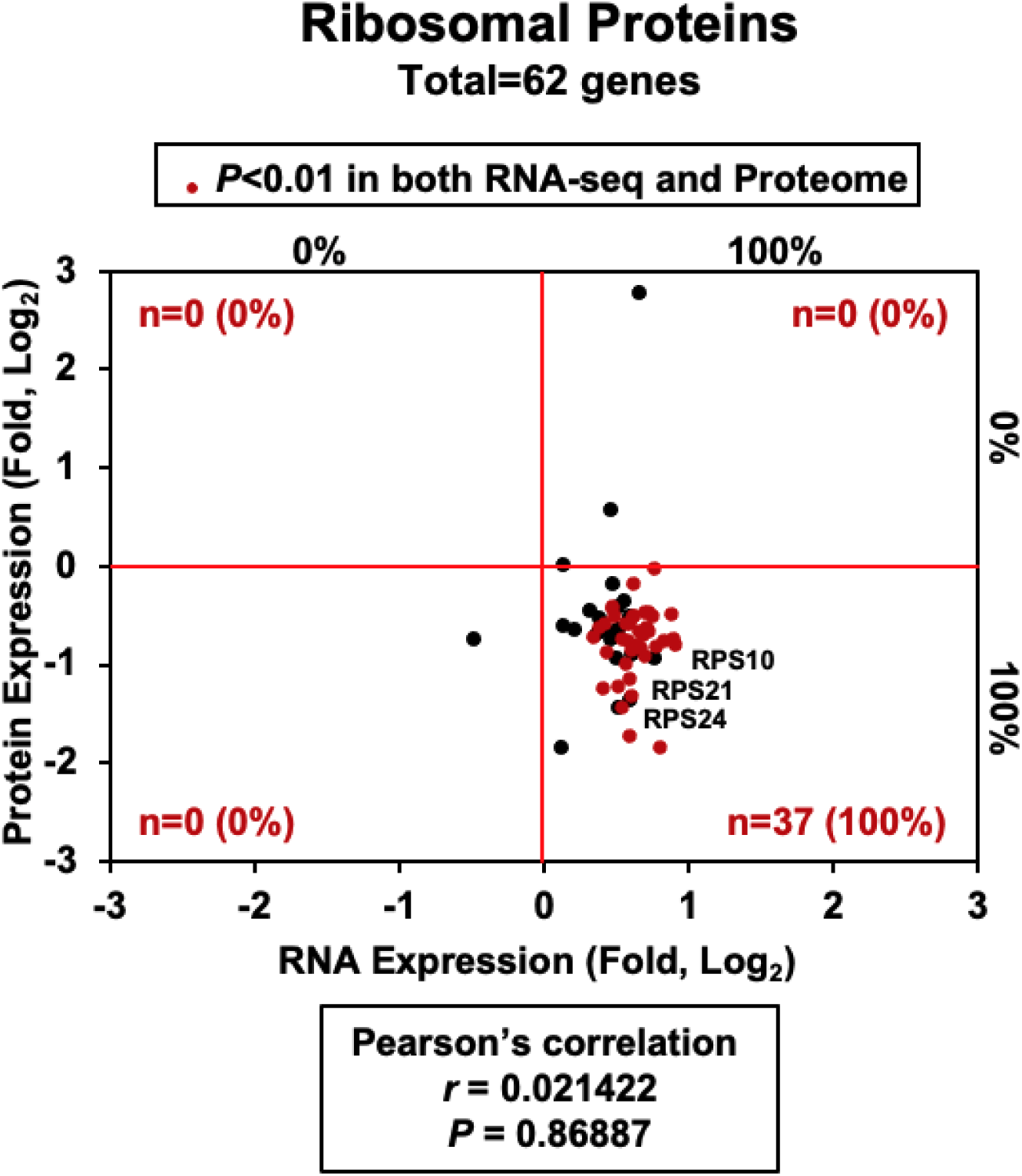
Post-Transcriptional Regulation of Ribosomal Proteins by *PPARα*. Integrative analysis of RNA and protein expression levels using ribosomal protein gene sets, with 62 genes overlapping the common dataset. The Pearson correlation coefficient was calculated to assess the concordance between RNA-seq and proteomics data, based on genes identified as statistically significant (P < 0.01).

**Supplemental Figure 5.**
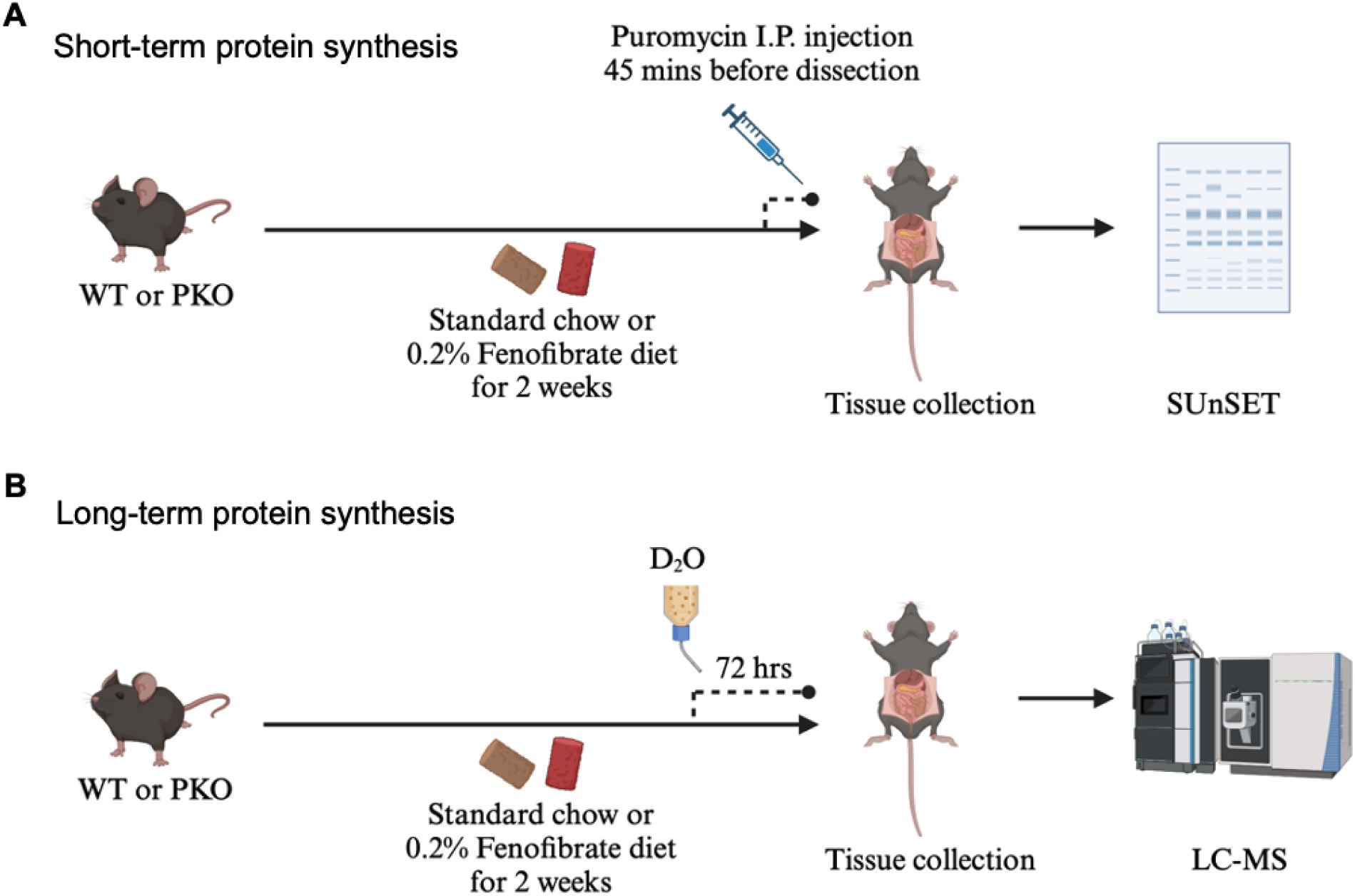
Overview of Experimental Approach to Measure Short- and Long-Turnover Protein Synthesis. (A) WT and PKO mice were fed either a standard chow or 0.2% fenofibrate-containing diet for two weeks. To assess short-turnover protein synthesis, mice were intraperitoneally injected with puromycin 45 minutes before sacrifice. Liver tissues were collected and analyzed using the SUnSET assay. (B) For long-turnover protein synthesis analysis, WT and PKO mice received 8% D₂O in drinking water for 72 hours during the final phase of the dietary intervention. Liver tissues were collected and analyzed via liquid chromatography–mass spectrometry (LC-MS).

**Supplemental Figure 6.**
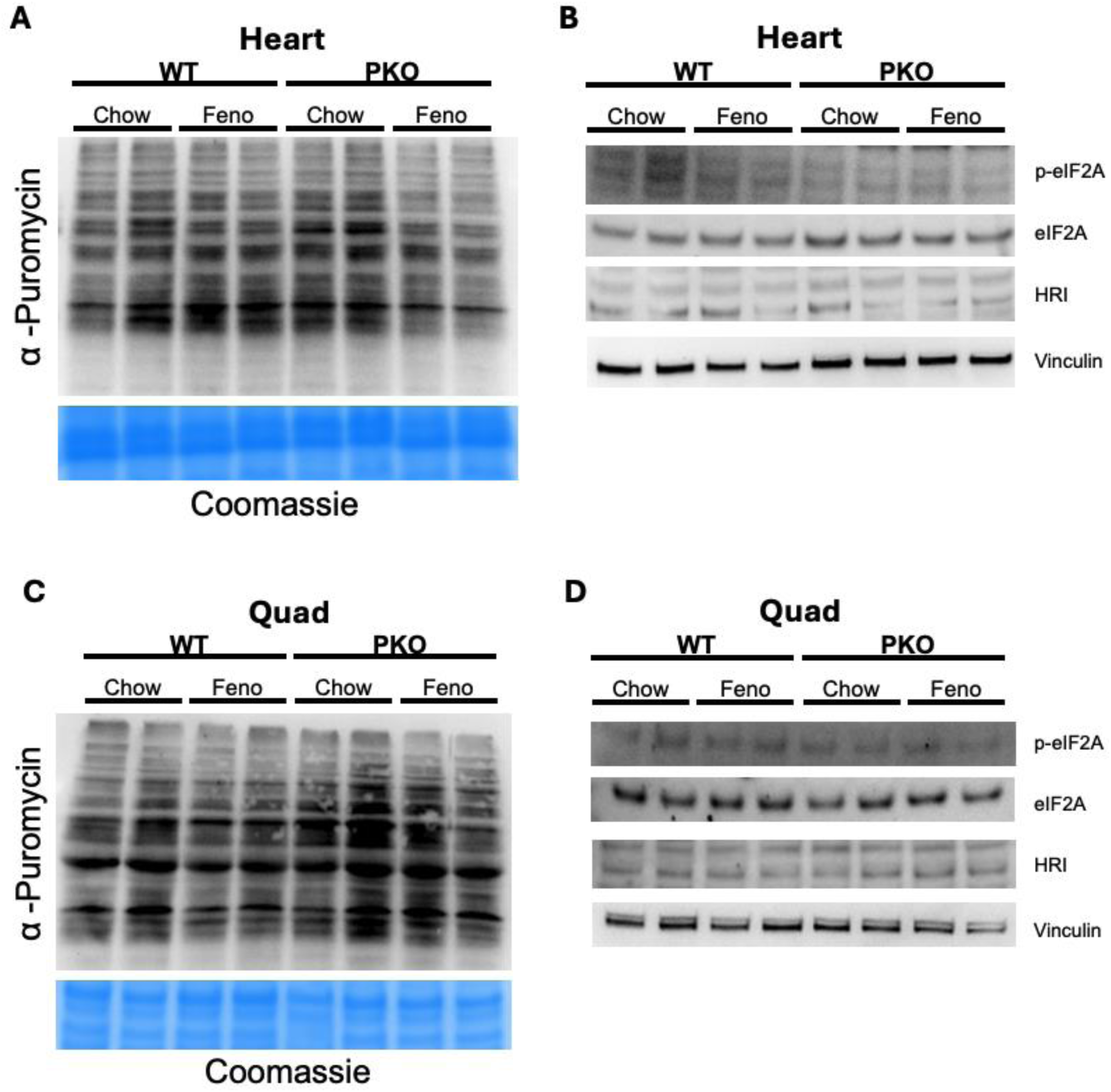
Tissue-Specific Effects of *PPARα*: No Repression of Protein Translation in Heart and Quadriceps. WT and PKO mice were fed either a normal chow diet or a chow diet containing 0.2% fenofibrate for two weeks. To assess short-turnover protein synthesis rates using the SUnSET assay, animals were intraperitoneally injected with puromycin at a dose of 40 nmol/g body weight. After 45 minutes, mice were sacrificed, and hearts and quads were harvested for further process. (A) Western blot–based SUnSET analysis of heart samples to assess short-turnover protein synthesis across the four experimental groups. Coomassie staining was used as a loading control (n = 2 per group). (B) Western blot analysis of phosphorylated and total forms of eIF2A and HRI in heart samples from the four experimental groups (n = 2 per group). Vinculin was used as a loading control. (C) Western blot–based SUnSET analysis of quad samples to assess short-turnover protein synthesis across the four experimental groups. Coomassie staining was used as a loading control (n = 2 per group). (D) Western blot analysis of phosphorylated and total forms of eIF2A and HRI in quad samples from the four experimental groups (n = 2 per group). Vinculin was used as a loading control.

